# A virtual cohort framework with applications to adoptive cell therapy in bladder cancer

**DOI:** 10.64898/2026.03.06.710135

**Authors:** Hannah G. Anderson, Sarah Bazargan, David J. Nusbaum, Michael A. Poch, Shari Pilon-Thomas, Katarzyna A. Rejniak

**Author notes:** Corresponding author (HGA), (KAR).

## Abstract

Even under the same treatment, responses can vary. Virtual cohorts build on an available, often limited, dataset can capture these differences and enable the discovery of treatment protocols that work well for a wide variety of individuals. In this paper, we refined current virtual cohort pipelines by improving data handling, ensuring the virtual cohort can be used to stratify individuals into treatment subgroups based on their data, and validating that the virtual cohort matches the observed data variability. To illustrate, we applied this pipeline to a murine data set of orthotopic bladder cancer treated with gemcitabine (Gem) and immunotherapy with OT-1 cells. We generated over 10,000 virtual mice that replicate the dynamics of three cell subpopulations in the tumor (cancer cells, T cells, and myeloid-derived suppressor cells) and data from four experimental cohorts (control, Gem, OT-1, and Gem+OT-1). We also provided a guide for using this pipeline for other treatments.

## 1 Introduction

The development of new therapies and treatment regimens is a multi-step process that includes in vitro experiments, preclinical animal studies, and four phases of clinical trials. In particular, animal studies and phase I clinical trials assess the feasibility, safety, and/or dosing of treatments using small treatment cohorts. Preclinical studies usually use 5-10 animals per treatment group, while phase I clinical studies are designed for 9-12 patients, with rigorous protection of trial participants, continuous monitoring of patients’ responses and side effects, and strict stopping rules if dose-limiting toxicities or adverse reactions are observed. For those small cohorts of animals or patients, often quite different responses are recorded, even under the same treatment. Some treated individuals may have more aggressive disease but respond well to treatment, while others have a slow-progressing disease but are unresponsive to treatment. To capture the variability of such responses while simultaneously providing a much larger group of simulated individuals (mice or patients) that faithfully reproduce the limited data, the concept of virtual clinical trials has been proposed^1–7^. Moreover, there is an ongoing trend to reduce the number of animals used in preclinical studies in favor of so-called New Approach Methodologies (NAMs), such as in vitro human-derived organoids, organ-on-a-chip technologies, and in silico computational models ^8,9^. Given this current need and context, we propose a pipeline to create a virtual cohort that expands a small experimental data set into a larger collection of virtual individuals, reproducing data faithfully and incorporating the data variability observed within the experimental cohort.

In the last 10 years, there have been several approaches detailing pipelines for generating a virtual cohort^1–7^ using clinical or animal data, although the development of in silico trials goes back further^10^. Chase et al.^1^ starts our conversation by asking vital questions necessary for building a good virtual cohort in the clinical setting. These questions pertain to physiological and clinical relevance of the model as well as treatment sensitivity. When constructing a model to address the problem, first ask the question: *is this model a good representation of the relevant biological processes for the problem?* Chase et al.^1^ also encourages the focus on clinical relevance by asking *are the model inputs and outputs relevant in guiding clinical care?* And, *is the model identifiable with respect to clinical data from the bedside?* Lastly, since the virtual cohort is developed for therapeutic testing, one should ask: *can we predict patient response to treatment input?* Chase et al.^1^ also included several ways to validate a virtual cohort, such as “cohort level before and after” to compare the variability of the virtual cohort to the experimental data and “cohort-level cross-validation” to evaluate the cohort’s ability to generalize to alternative protocols.

A year after the work by Chase et al.^1^, Viceconti et al.^2^ proposed a process for developing a virtual cohort that first considers the model’s context of use in the regulatory process (Step 1), thus addressing the biological and clinical relevance questions put forth by Chase et al.^1^. Then, after code and model verification as a quality check (Step 2) and estimating parameters (Step 3), Viceconti et al.^2^ suggested performing model sensitivity to determine parameters that likely cause model (and hence cohort) variability as well as uncertainty quantification (Step 4). Uncertainty quantification (e.g., parameter identifiability) addresses the questions regarding model identifiability with data and predicting patient response to treatment. Lastly, the virtual cohort pipeline ended with cohort validation while keeping in mind the pre-defined context of use (Step 5)—although specific validation ideas were not suggested like Chase et al.^1^—and checking (Step 6) that the model meets technical standards set forth by the American Society of Mechanical Engineering^11^. Around the same time as Viceconti et al.^2^, Sinisi et al.^3^ proposed a framework for producing virtual patients for a non-identifiable model that uses Statistical Model Checking and hypothesis testing to obtain a complete spectrum of possible patients while using an informed sampling policy to maintain that each patient is distinguishable in the context of parameter interdependence.

Then, a few years later, Arsene et al.^4^ revisited the pipeline proposed by Viceconti et al.^2^, but rearranged the order by grouping inter/intra patient variability and model calibration into the same step (as virtual population design), and then grouped model validation, verification, and uncertainty quantification under the context of regulatory guidance^11^ into the next step. They defined validation as a check that the model assumptions are correct and the uncertainties and sensitivities are understood, while verification confirmed that the mathematical model is correctly coded and accurately solved. Arsene et al.^4^ also mentioned curating a knowledge model of relevant biomedical knowledge between Steps 1 and 2 of Viceconti et al.^2^—a step that was implicitly included in Viceconti et al.^2^ but by explicitly including it in the pipepline, Arsene et al.^4^ gave more advice as to how to create a relevant model.

Craig et al.^5^ streamlined this pipeline into a practical guide for creating virtual clinical trials with the following steps: (1) create model informed by available data, (2) parameterize model, (3) conduct sensitivity and identifiability analysis, (4) create virtual cohort, (5) conduct virtual clinical trial to answer the question, and then (6) revise as needed. Craig et al.^5^ gave more hands-on guidance than previous papers by explicitly laying out sensitivity and identifiabilty analysis definitions and methods. Gevertz and Wares^6^ had the same pipeline as Craig et al.^5^ except gave explicit instructions as to how to determine plausible patients by using either “accept-or-reject” or “accept-or-perturb” methods^12^. Kleeberger’s^7^ review of virtual cohort generation also had the same pipeline as Craig et al.^5^ but furthered discussion by considering alternative methods for creating virtual patients (agent-based models, AI/ML, digital twins, biosimulation/statistical methods), although some of these methods are difficult to scale to a virtual cohort.

We propose an improvement to this established pipeline by performing a priori structural identifiability analysis to ensure proper data handling before parameterization given the model’s structure. To illustrate the impact of skipping this step, an oncology model can fit tumor volume data well while lacking the ability to accurately capture the dynamics of modelled cell types within the tumor. This has implications for treatment response predictions, especially to immunotherapies. Craig et al.^5^ mentioned a posteriori structural identifiability analysis, which would occur after the initial parameter estimation for population-level parameters. However, it may be that the model’s structure or the data type used was not suitable for the population-level parameter estimation. A posteriori methods use an actual data set to determine parameters that cannot be identified, whereas a priori methods only use the model structure to determine if a *type* of data can identify model parameters^13^. Since a posteriori methods (such as identifying flat profile likelihoods) use an actual data set, if a parameter is not structurally identifiable, the issue can lie with aspects about the experimental data set, the model’s structure, or the data type. This makes it difficult to know how to resolve this issue. This is in contrast with a priori methods, where if a parameter is not determined to be structurally identifiable, the issue is resolved either by altering the model’s structure or by changing the data type. Thus, we suggest the use of a priori methods because they give clearer guidance in the case that the model is not structurally identifiable. Further, we suggest global structural identifiability rather than local since global is a stronger conclusion that does not depend on suggested parameter ranges or an initial guess.

We also further refined the order of the pipeline by suggesting that, after population-level parameter estimation, one should first identify several practically identifiable parameter subsets based on longitudinal data and then perform a sensitivity analysis to determine which of these subsets cause the most variability in the model output. Using this order increases the variability produced by the cohort while maintaining that the generating subset can be used in the future to create a digital twin based fitting the practically identifiable subset to an individual’s longitudinal data. These digital twins improve future patient (or animal) stratification given the circumstance where different subgroups within the virtual cohort respond better to different treatment regimens. Since a practically identifiable parameter set differentiates these subgroups, patients (or animals) can be stratified into these subgroups using their digital twin in order to maximize treatment efficacy for that individual. We also note that, after practical identifiability and sensitivity analysis, one should check that the generating parameter subset exhibits variability experimentally to ensure the feature causing variability between virtual patients also causes variability between experimental.

Additionally, our goal was to develop a virtual cohort that can be applied to data from clinical trials or animal experiments with multiple arms including controls, monotherapies, and combination therapies. Using the “accept-or-reject” method from Gevertz and Wares^6^, a parameter set is accepted into the virtual cohort as a virtual patient (or subject) if its simulations are within a realistic distance (e.g. 2 or 3 standard deviations) from an average individual under treatment at each data collection time point. We expand this to include multiple arms by suggesting that each virtual individual be simulated under the different treatment arms and accepted only if simulations are reasonable for all time points and all treatment cohorts.

Lastly, our pipeline revisits the validation methods suggested by Chase et al.^1^ to compare virtual and experimental cohorts. If one wants to perform “cohort-level cross-validation” to evaluate the virtual cohort’s generalizability to other regimens, a treatment cohort can be left out of the “accept-or-reject” method for this analysis. For both this analysis and “cohort-level before and after”, we suggest an explicit method—the Kolmogorov-Smirnov test—to compare the distributions of the virtual to experimental cohorts. Ensuring that the representation observed in the virtual cohort is realistic is essential as a skewed representation will interfere with future testing of treatment regimen robustness on the cohort. For instance, the virtual cohort could overpredict response to a regimen largely because it overrepresents a certain patient (or animal) subgroup. Since testing treatment robustness is often a main reason for developing virtual cohorts, these validation methods are essential to obtain accurate predictions in line with experimental variability.

As a basis for applying this pipeline, we used experimental murine data on orthotopic bladder cancer treated with combined chemotherapy and adoptive cell therapy (ACT) using T cells, published by our group ^14^. In these experiments, mice were intravesically (inV) implanted with MB49 bladder tumors expressing ovalbumin (MB49-OVA) and then treated with inV injections of gemcitabine (Gem) at day 10 post-implantation and inV OT-1 T cells at day 14. The low-dose Gem was used to remove immunosuppressive cells, such as myeloid-derived suppressor cells (MDSCs) ^15^. OT-1 T cells recognize the ovalbumin (OVA) antigen ^16^, thus this murine model serves as a surrogate for adoptive cell therapy with tumor-infiltrating lymphocytes (TILs) in patients ^17^. The experiments described in ^14^ comprise four arms: control, Gem and OT-1 monotherapies, and the combined Gem+OT-1 treatment. Flow cytometry data of CD8^+^ T cells and MDSCs collected on day 14 confirmed that Gem preconditioned the tumor-immune microenvironment for OT-1 treatment by decreasing the immunosuppressive MDSC population (^14^, Fig 4). Further, ultrasound data collected at days 6, 9, 13, 16, and 20 indicated that Gem+OT-1 combination treatment showed a statistically significant decrease in the size of the tumor versus all other treatment cohorts tested: untreated, Gem monotherapy, and OT-1 monotherapy treated mice (^14^, Fig. 6C). However, this improved response was also coupled with variability in treatment response in the murine population (^14^, Fig. 6B), thus there was a rationale to develop a virtual murine cohort that reproduces experimental dynamics and captures the variable tumor response and growth. Such virtual cohorts open the possibility to test alternative treatment strategies to identify robust regimens that work better for more individuals. However, these improved treatment protocols were out of the scope of this paper, so we concentrated on the virtual cohort pipeline.

Our paper includes a description of the pipeline (Section 2.1), the mathematical model (Section 2.2), structural identifiability analysis using the differential algebra approach (Section 2.3), parameter estimation (Section 2.4), practical identifiability analysis using profile likelihoods with ultrasound data (Section 2.5), and sensitivity analysis with the eFAST method to determine the most sensitive practically identifiable parameter subset (Section 2.6), the “accept-or-reject” method is used to create the virtual murine cohort (Section 2.7), which is validated in Section 2.8. We finish by presenting a guide for modelers who want to use the pipeline for their own problem (Section 2.9). Our paper concludes with a discussion of our virtual cohort and future directions for treatment optimization, robustness, and subject stratification for personalized therapy (Section 3). All methods are described in more details in Section 4.

## 2 Results

### 2.1 Virtual cohort pipeline

The flowchart of the virtual cohort pipeline is presented in Fig 1 and briefly described below.

**Fig 1.**
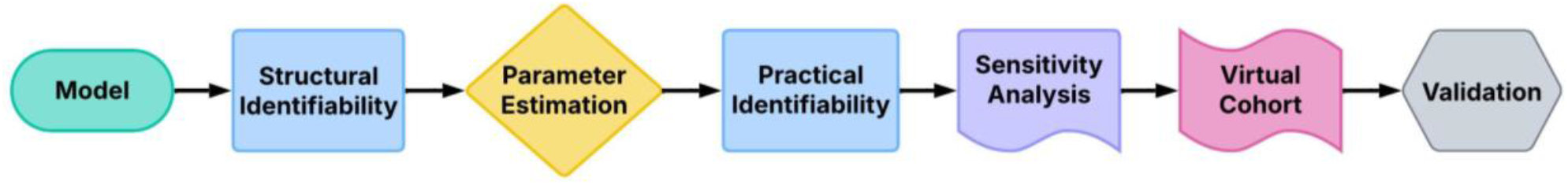
Virtual cohort pipeline. Pipeline used in the creation of our virtual cohort.

Our virtual cohort pipeline consists of the following 7 steps:

1. **Model:** First, we develop a fit-for-purpose ordinary differential equations (ODE) model containing cell populations and therapies based on knowledge of the biology, mechanisms of the treatment(s), and the available data.
2. **Structural Identifiability**: Then, we perform a priori structural identifiability analysis to determine if the available data is sufficient to identify model parameters given the model’s structure. Our analysis uses the differential algebra approach. This is an important step since some data may be unsuited to identify the subpopulation dynamics of the model, especially if the model contains several cellular components.
3. **Parameter Estimation:** Results from structural identifiability analysis then inform our use of data in parameter estimation.
4. **Practical Identifiability:** Afterwards, we perform practical identifiability analysis using profile likelihoods to find subsets of parameters that can be identified with available longitudinal data. For data that can be obtained in minimally invasive way (i.e., ultrasound or radiologic images), this enables the future creation of digital twins, which can be used to stratify subjects for adaptive therapy.
5. **Sensitivity analysis:** Next, sensitivity analysis discerns the practically identifiable subset that is best at capturing variability in the model output and thus may be better suited to capture variability in the virtual cohort. We use the eFAST method here.
6. **Virtual Cohort Generation:** After checking that the identified parameter set also exhibits variability experimentally, we use this set to generate the virtual cohort using the “accept-or-reject” method ^6^.
7. **Validation:** Lastly, we conclude the pipeline with virtual cohort validation using the “cohort-level before and after” method ^1^ for data reproduction, “cohort-level cross-validation” ^1^ for generalizability, and demonstrate that individual mice growth curves are well represented in the virtual cohort.

This virtual cohort pipeline was applied to murine data of bladder cancer under treatment with Gem and OT-1 cells from our group ^14^. However, this pipeline could also be used for data from other mice experiments or for patient data collected in clinic.

### 2.2 Mathematical model for Gem + OT-1 combination therapy

Based on our murine data ^14^, we developed a fit-for-purpose 4-equation ODE model of cancer cells (*C*), CD8^+^ T cells (*T*), and MDSCs (*M*) that incorporates treatment with OT-1 cells (*T*), and Gem (*G*). All cell subpopulations were modelled in terms of their volume (*mm*^3^), whereas Gem was in terms of concentration (*μM*). Fig 2 showcases the flowchart of tumor-immune-Gem-OT-1 interactions described by the model.

**Fig 2.**
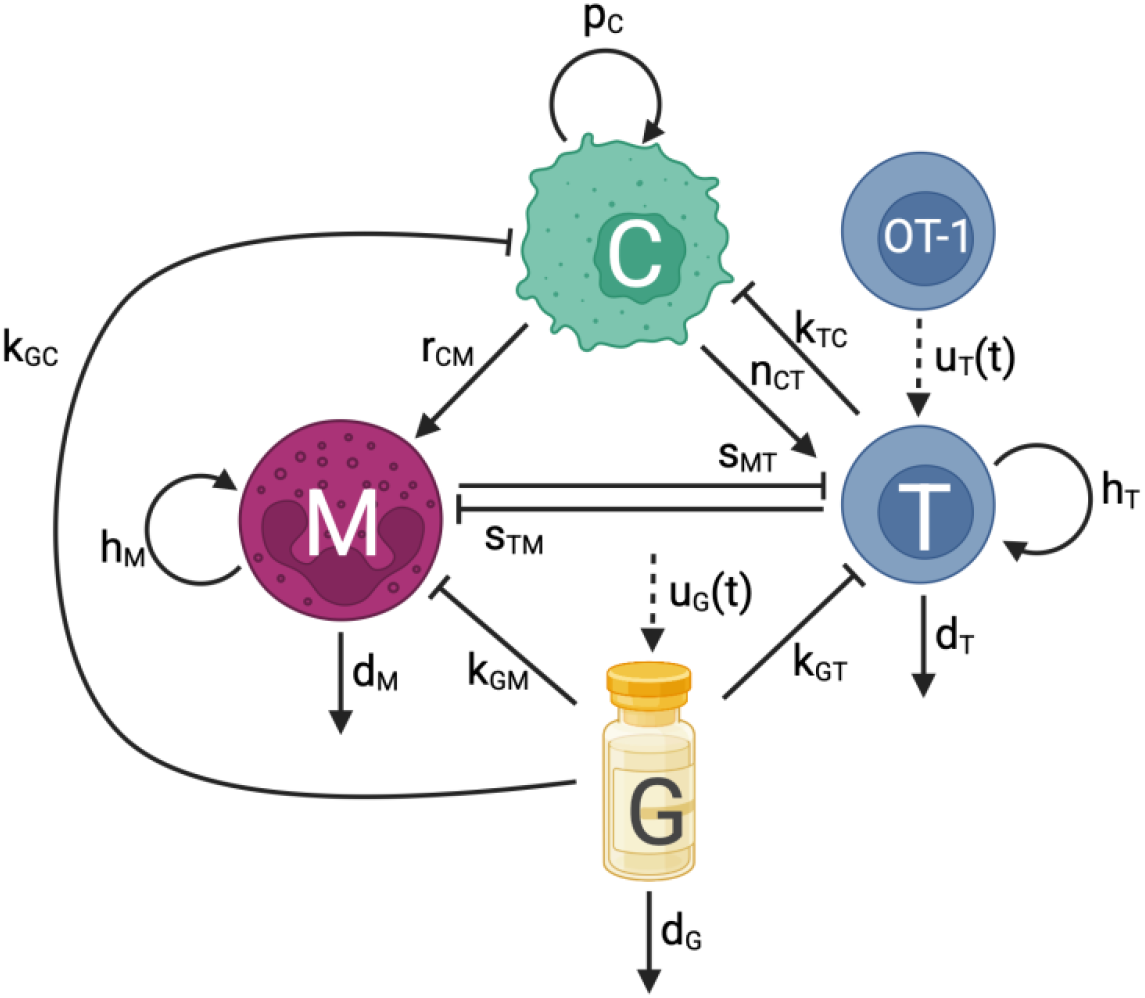
Flowchart of tumor-immune-Gem dynamics. Flowchart displaying the interactions of cancer cells (*C*), CD8+ T cells (*T*), and MDSCs (*M*) under treatment with OT-1 cells and Gem (*G*). Sharp solid arrows represent proliferation, recruitment, or removal. Blocked arrows represent killing or suppression. Dashed arrows represent treatment injections. Flowchart was created with BioRender.com.

#### Model equations

***cancer cells*** (***C***):

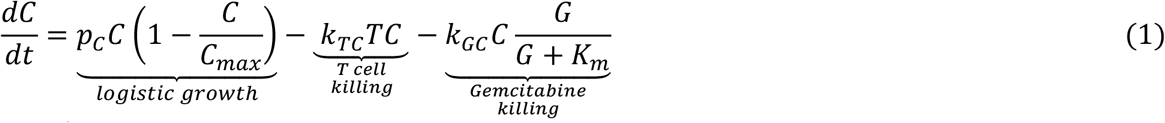

***CD*8**^+^ ***T cells*** (***T***):

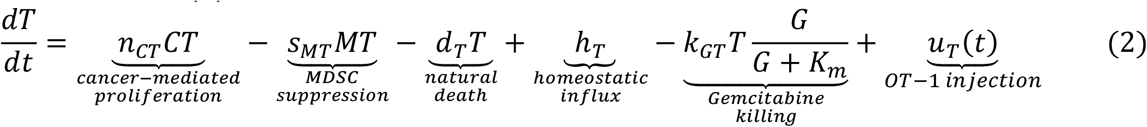

***Myeloid***™***derived suppressor cells*** (***MDSCs***) (***M***):

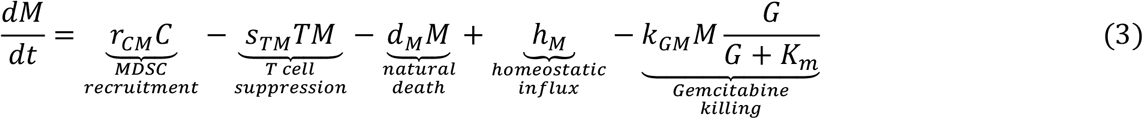

***Gemcitabine*** (***G***):

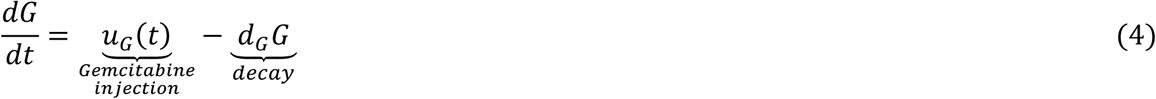

In equation 1, we assumed that cancer (*C*) grows logistically at a rate of *p*_*C*_ with a tumor carrying capacity of *C*_*max*_. T cells kill cancer cells at a rate of *k*_*TC*_. Since ^18^ found that Gem uptake is saturable and follows Michaelis-Menten kinetics, Gem killing of cancer was modelled using the Michaelis-Menten equation, where the Michaelis constant, *K*_*m*_, is the Gem concentration at which the transporter uptake rate is at half its maximum velocity, and the maximal kill rate is *k*_*GC*_. All cell types were assumed to have the same Michaelis constant, *K*_*m*_, since the same transporter protein, equilibrative nucleoside transporter 1 (ENT1), is used for Gem uptake by bladder cancer cells and T cells ^18–20^. Further information regarding the Michaelis constant can be found in Supplementary Section S1.

In equation 2, T cells (*T*) undergo expansion at a rate of *n*_*CT*_ due to the presence of a tumor, as ^21^ experimentally noticed an increase of proliferating CD8^+^ T cells in the center of the tumor versus the invasive margin for several tumor types including bladder cancer. T cells experience immune suppression by MDSCs at a rate of *s*_*MT*_^22^. They also die naturally at a rate of *d*_*T*_. Since histology data from non-tumor bearing mice showed a T cell population within the bladder, we included a term for the homeostatic influx of T cells, *h*_*T*_, regardless of the presence of a tumor, where *h*_*T*_ = (*s*_*MT*_*M*_0_ + *d*_*T*_)*T*_0_ according to our analysis of the tumor-free equilibrium, (0, *T*_0_, *M*_0_), in Theorem A of Supplementary Section S2. Similar to the cancer equation, Gem saturates according to Michaelis-Menten dynamics with a Michaelis constant of *K*_*m*_ and a T cell kill rate of *k*_*GT*_. OT-1 treatment administration was modelled by the time-dependent function, *u*_*T*_(*t*).

In equation 3, MDSCs (*M*) are recruited by the cancer cells to the tumor site at a rate of *r*_*CM*_. MDSC recruitment, which can occur through mechanisms such as CXCL2/MIF-CXCR2 signaling in bladder cancer, is associated with a poor prognosis ^23,24^. T cells also cause a decrease in MDSCs at a rate of *s*_*TM*_. This term was motivated by the decrease in MDSCs seen in the tumor histology of mice treated with Gem combined with OT-1 (14.7% Ly6G MDSCs in tumor) compared to mice treated only with Gem (2.4%) at day 17 post-implantation (Fig 3). This data indicated that T cells may have some impact on decreasing the number of MDSCs. MDSCs die naturally at a rate of *d*_*M*_. Because MDSCs were present in the normal bladder tissue (as per our murine data), the MDSC homeostatic influx rate, *h*_*M*_, was included in the model and can be substituted by *h*_*M*_ = (*s*_*TM*_*T*_0_ + *d*_*M*_)*M*_0_ (Theorem A of Supplementary Section S2). Gem saturates with a Michaelis constant of *K*_*m*_ and kills MDSCs at a maximal rate of *k*_*GM*_.

**Fig 3.**
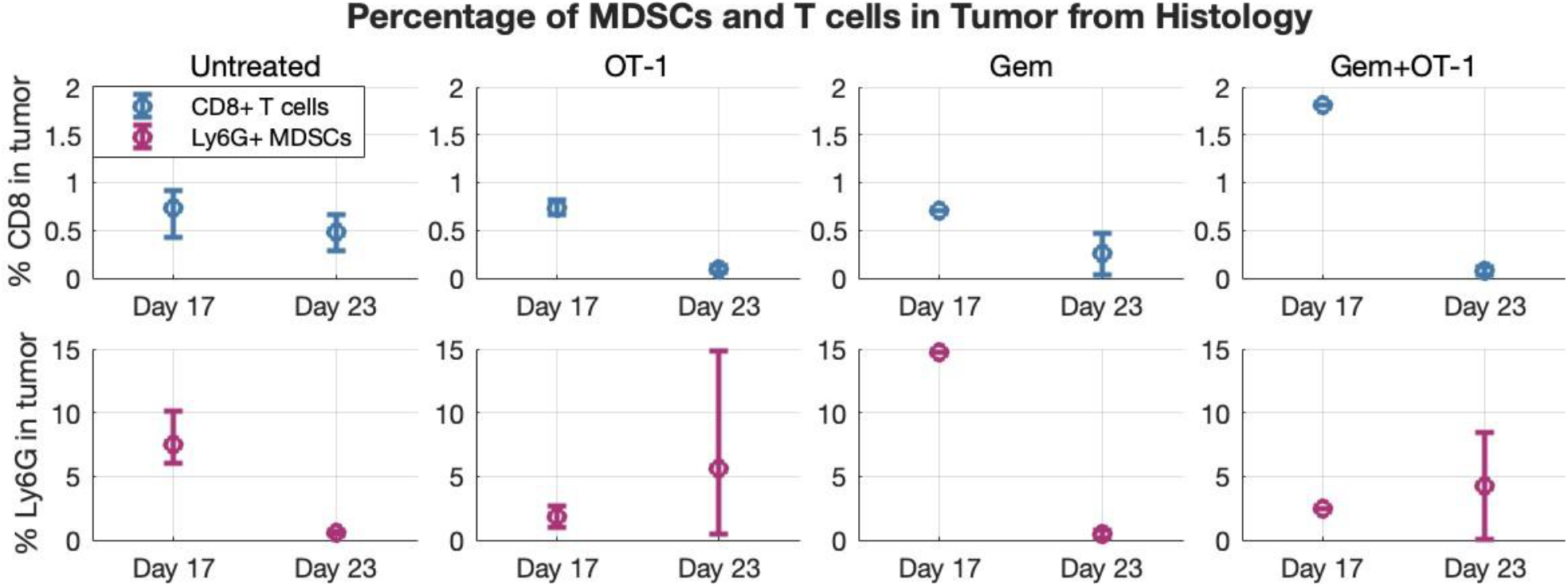
CD8^+^ T cell and MDSC histology data. Percentage of the tumor (in terms of area on the histology slide) that is Ly6G^+^ MDSC and CD8^+^ T cells for each of the four treatment groups on days 17 and 23 post-tumor implantation. Error bars represent the minimum and maximum of the data point compared to the mean (circles).

In the 4^th^ and final equation, Gem (*G*) is injected according to schedule, *u*_*G*_(*t*), and then decays at a rate of *d*_*G*_. Supplementary Table S1 contains parameter values, ranges, descriptions, and the best fit to our data.

### 2.3 Structural identifiability analysis

Given that the majority of our data were ultrasounds capturing the total volume of mice bladder tumors (^14^, Fig. 6C), we wanted to answer the question: *can ultrasound data be used to identify model parameters?* If not, *what type of data should be used?*

By definition, a model is globally structurally identifiable with respect to a data type if each parameter has a unique value consistent with the observed model outputs. In the following section, we used the differential algebra approach (Section 4.1.1) developed by ^25^ to show that the model is not globally structurally identifiable with respect to data on the total tumor volume (Theorem 1), but it is with respect to data on each cell subpopulation and Gem (Theorem 2).

#### 2.3.1 Structural identifiability theorems

For Theorem 1, we used the simpler version of the model without any (Gem or OT-1) treatment:

***cancer cells*** (***C***):

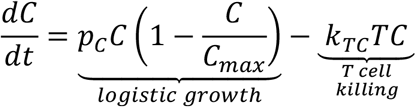

***T cells*** (***T***):

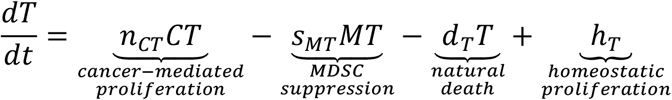

***Myeloid***™***derived suppressor cells*** (***MDSCs***) (***M***):

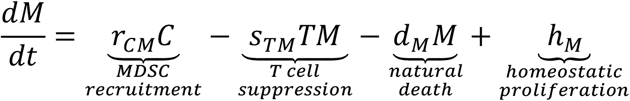

If the treatment-free model is not globally structurally identifiable, then the full version of the model including treatment (and thus more parameters) is also not globally structurally identifiable.

##### Theorem 1.

The treatment-free model is not globally structurally identifiable with respect to total volume data, *C* + *T* + *M*.

**Proof**.

According to our treatment-free model, the change in total tumor volume over time is

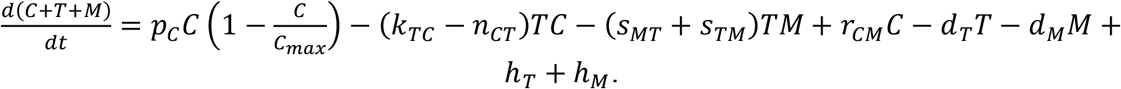

So, our input-output relation is

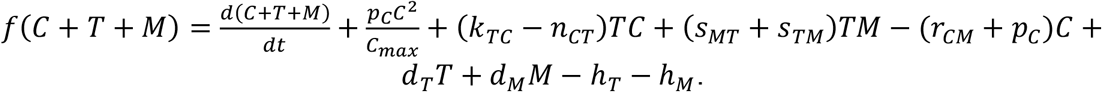

Now, according to the differential algebra approach ^25^, the identifiable parameter combinations are the coefficients of *C* + *T* + *M* and its derivatives in the input-output relation. Since it is difficult (if not impossible) to simplify the right-hand side to be solely in terms of *C* + *T* + *M* and its derivatives, we took a different approach.

We considered the first equation in this proof to be our model in terms of the change of the total tumor volume over time, and we assume that we have data on *C, T*, and *M*. According to our input-output relation, the identifiable parameter combinations are:

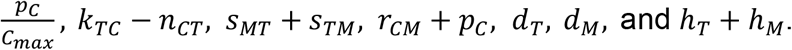

Since we have 7 combinations but 11 parameters, the parameters are not independent and their values cannot be deduced from these combinations. Therefore, the model in terms of total tumor volume (first equation in proof) is not structurally identifiable with respect to *C, T*, and *M*, and thus, it cannot be structurally identifiable with respect to *C* + *T* + *M* data.

From Theorem 1, we concluded that ultrasound data is insufficient to fit all parameters and thus capture the subpopulation dynamics of the treatment-free model. Since this is true for the model without treatment, it is insufficient for any of the treatment cohorts.

In addition to ultrasound data, flow cytometry measurements and histology images were collected during experiments described in ^14^. Flow cytometry data was collected from 12-13 mice on day 14 for the untreated and Gem-treated groups (^14^, Fig. 4), and histology data was collected for 1-3 mice per treatment group on day 17 and 23 and converted to ratios of T cell and MDSC area in comparison to the area of the tumor on the histology slide (Fig 3).

**Fig 4.**
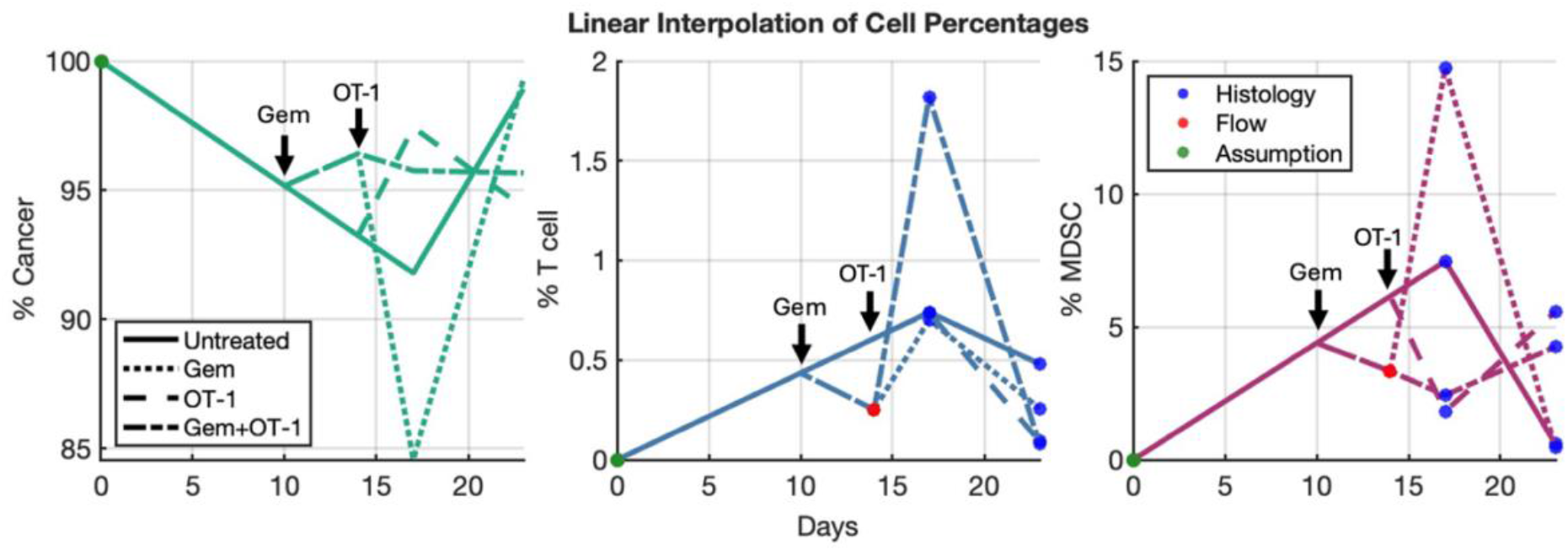
Linear interpolation of cell subpopulation data. Linear interpolation of the cell percentages in the tumor microenvironment for the four treatment cohorts using histology and flow cytometry data.

Using the doses of injected OT-1 and Gem as well as the Gem decay time (3 days), we checked if having data on each cell type and Gem is sufficient to determine parameters for the full model in Theorem 2. If the full model with treatment is globally structurally identifiable with respect to this data, then we can use this result to identify parameters for any of the experimental cohorts.

##### Theorem 2.

*The full model is globally structurally identifiable with respect to data on C, T, M, and G*.

**Proof**.

Our input-output relations are formed by rearranging the four ODEs in equations (1)–(4) to equal to 0 and then multiplying by any denominators. This forms the following polynomials in terms of *C, T, M, and G* and their derivatives:

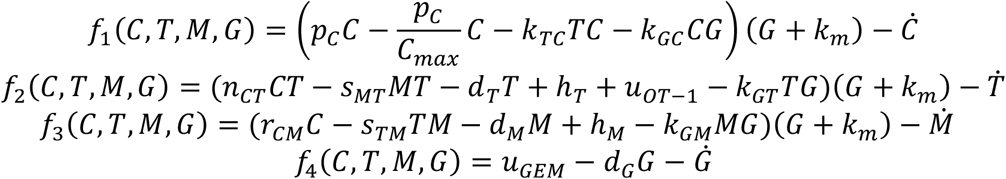

The set of coefficients from *f*_1_, *f*_2_, *f*_3_, and *f*_4_ form identifiable combinations of parameters. Specifically, using the following coefficients:

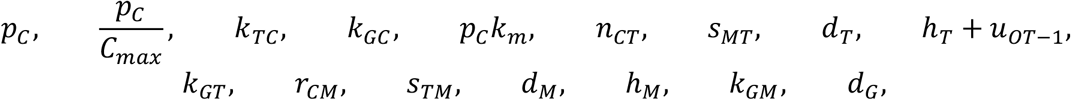

we can analytically express all 16 parameters; note, that the OT-1 administration dose, *u*_*OT*−1_is known. Therefore, the model is globally structurally identifiable with respect to data on *C, T, M*, and *G*.

#### 2.3.2 Application of structural identifiability result

We showed that our model is not structurally identifiable with respect to total tumor volume data, but it is with respect to volume data on each cell subpopulation and Gem. The necessary data for Gem—administration dose and decay—are know from experiments. Therefore, we only needed to determine volumes for cancer, T cells, and MDSC populations over time within the tumor microenvironment.

Using the average histology (Fig 3) and flow cytometry (^14^, Fig 4) data at 3 different time points, we performed a linear interpolation to determine the percentage of the tumor that is CD8^+^ T cells, Ly6G^+^ MDSCs, and cancer cells (Fig 4). We assumed that the tumor was 100% cancer on implantation day, the linear interpolation of untreated histology data defined the cell percentages until treatment started at day 10, and the Gem to untreated ratio from flow cytometry was used to distinguish between Gem-treated and non-Gem-treated cohorts at day 14. We multiplied the ultrasound data by these percentages to obtain volumes for each cell type over time (Fig 5). Although this was not an exact measurement of cell subpopulation volumes, it gave an approximation with which to identify model parameters.

**Fig 5.**
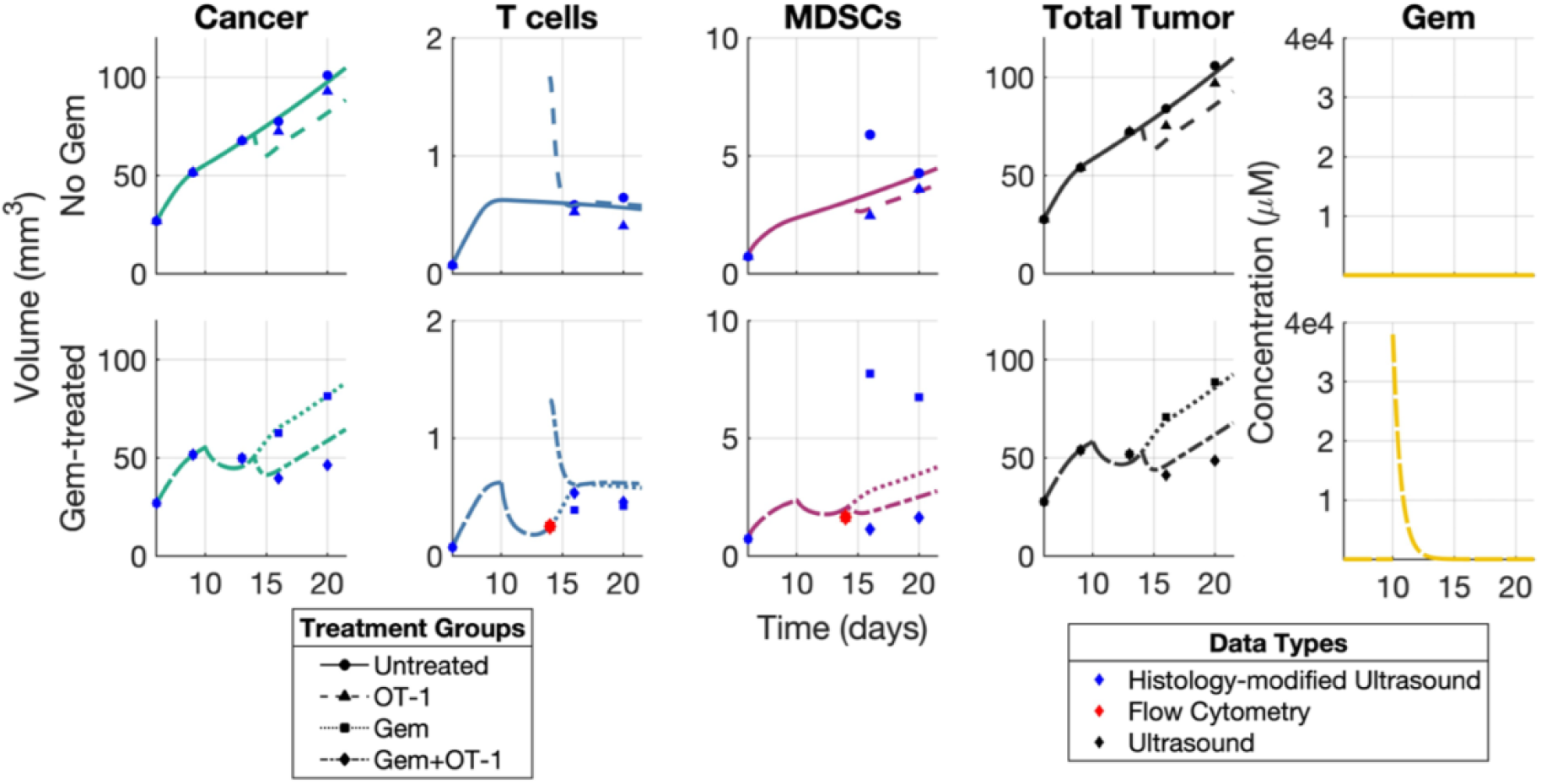
Numerical simulations after parameter estimation. Subpopulation volume curves generated for the hierarchically fitted parameters for subpopulation data from (top) untreated and OT-1-treated mice cohorts, and (bottom) Gem-treated and Gem+OT-1-treated mice cohorts.

### 2.4 Parameter estimation

We fixed the following parameters based on known values from literature or data (Supplementary Table S1): *d*_*T*_, *r*_*CM*_, *d*_*M*_, *d*_*G*_, and *K*_*M*_, and then fit the remaining parameters to subpopulation data using the gradient descent method (Fig 5). The parameters were fit hierarchically by first fitting *p*_*C*_, *C*_*max*_, *k*_*TC*_, *n*_*TC*_, *s*_*MT*_, *T*_0_, *s*_*TM*_ and *M*_0_ to the two non-Gem-treated cohorts and then fixing those parameters and fitting *k*_*GC*_, *k*_*GT*_ and *k*_*GM*_ to the two Gem-treated cohorts. Since the model with and without OT-1 treatment used the same parameters, each of the hierarchical steps produced the best fitting parameter set based on combined error of two cohorts. To be more precise, we calculated the relative error for each data point and then summed this across the three cell populations and the two cohorts being estimated:

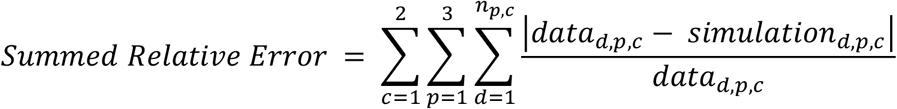

where *c* spans the cohorts tested, *p* spans the cell populations, *d* spans the number of data points (*n*_*p,c*_) for that cell population, *p*, and cohort, *c*.

To try to avoid gradient descent converging to a local minima, we sampled the parameter space using Latin Hypercube Sampling and then used the parameter set with the lowest error as our initial guess. Our initial condition at day 6 was 26.85 *mm*^3^ cancer cells, 0.07217 *mm*^3^ CD8+ T cells, and 0.7288 *mm*^3^ MDSCs, according to the linear interpolation of the three cell types (Fig 4) multiplied by the ultrasound mean at day 6. At day 10 post-implantation, Gem-treated cohorts received 3.8 × 10^4^ *μM* Gem. Then, on day 14, OT-1-treated cohorts received 1.0695 *mm*^3^ of OT-1. Since histology data for T cells and MDSCs was collected at days 17 and 23, we used the histology linear interpolation (Fig 4) to modify ultrasound volumes for T cells and MDSCs at day 16 and 20 as these days were closer to the histology collection days. By contrast, histology-modified ultrasounds for cancer cells were used at each ultrasound day since we assumed that the cancer cell dynamics more closely follow those of the total tumor over time.

Numerical simulations of the parameter set obtained under the four different regimens is plotted alongside data in Fig 5. More information about the fitted parameter set, parameter values and ranges obtained from literature, and a units conversion of the treatment injections can be found in Supplementary Section S1.

### 2.5 Practical Identifiability Analysis

Since the longitudinal data was collected from ultrasound measurements, the sampling and variability of our virtual cohort was dependent on the variability seen in our ultrasound data. Therefore, we wanted our generating parameter subset to be identifiable with respect to ultrasound data. To find this subset, we performed practical identifiability using the profile likelihood method (Section 4.2), which is the most common and accurate method ^26^.

We found that the profile likelihoods (Fig 6) for the tumor growth rate (*p*_*C*_), the homeostatic T cell population (*T*_0_), and the MDSC kill rate by Gem (*k*_*GM*_) exhibit a distinct local minimum showing that a unique value for each parameter explains the data set. Thus, these parameters are practically identifiable with respect to ultrasound data. Since different combinations of parameters can be practically identifiable, we continued testing and found four additional parameter combinations that are identifiable (Fig 6): *p*_*C*_, *r*_*CM*_ (MDSC recruitment rate), and *k*_*GM*_ (Set 2); *k*_*TC*_ (T cell kill rate of cancer cells), *r*_*CM*_, and *k*_*GM*_ (Set 3); *n*_*CT*_ (cancer-mediated proliferation of T cells), *r*_*CM*_, and *k*_*GM*_ (Set 4); *r*_*CM*_, *d*_*M*_ (MDSC death rate), and *k*_*GM*_ (Set 5). These results were consistent when we used the average ultrasound data from each of the four experimental cohorts: untreated, Gem and OT-1 monotherapies, and Gem+OT-1 combination therapy. Moreover, since non-invasive longitudinal data can uniquely identify these parameter subsets, the subsets can be used in the future to generate “digital twins” for experimental mice and stratify subjects for therapy.

**Fig 6.**
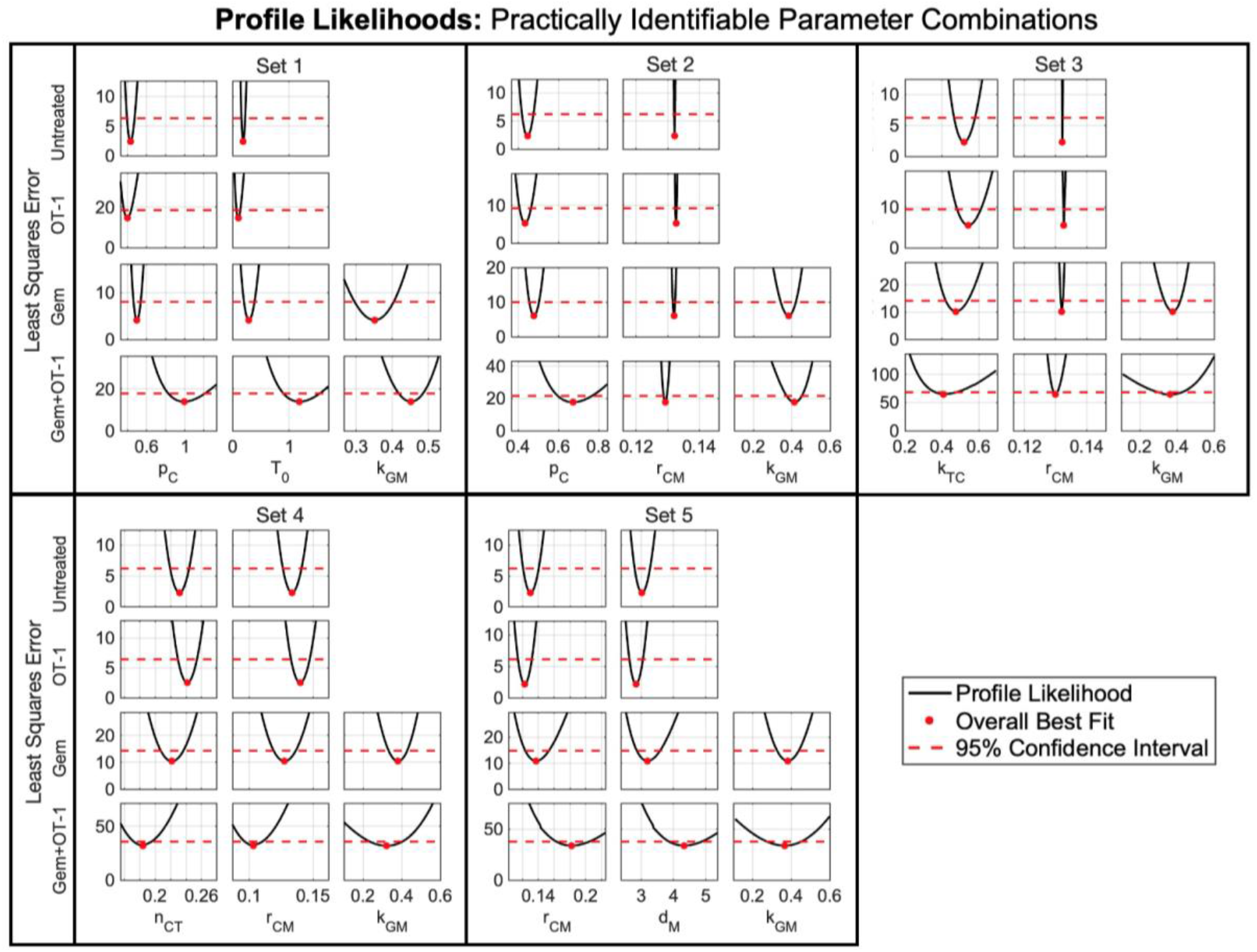
Practically identifiable parameter combinations. Profile likelihoods (black solid line) of the five sets of practically identifiable combinations of parameters (**Set 1**: *p*_*C*_, *T*_0_, and *k*_*GM*_; **Set 2**: *p*_*C*_, *r*_*CM*_, and *k*_*GM*_; **Set 3**: *k*_*TC*_, *r*_*CM*_, and *k*_*GM*_; **Set 4**: *n*_*CT*_, *r*_*CM*_, and *k*_*GM*_; **Set 5**: *r*_*CM*_, *d*_*M*_, and *k*_*GM*_) along with the 95% confidence threshold (red dashed line) for the four experimental treatment cohorts. Since profile likelihoods show a distinct local minimum, these five combinations of parameters are practically identifiable with respect to ultrasound data.

### 2.6 Sensitivity Analysis

To generate the virtual cohort, we wanted to select the parameter subset from Section 2.5 that most varied the model output (i.e., the subset that the model was most sensitive to), as this subset would better capture the experimental variability during sampling. Using the extended Fourier amplitude sensitivity test (eFAST) described in Section 4.3, we determined the model’s sensitivity to parameters (Fig 7). Since the five sets of practically identifiable parameters each included the same Gem-related parameter, *k*_*GM*_, we only compared the model’s sensitivity to parameters not related to Gem treatment.

**Fig 7.**
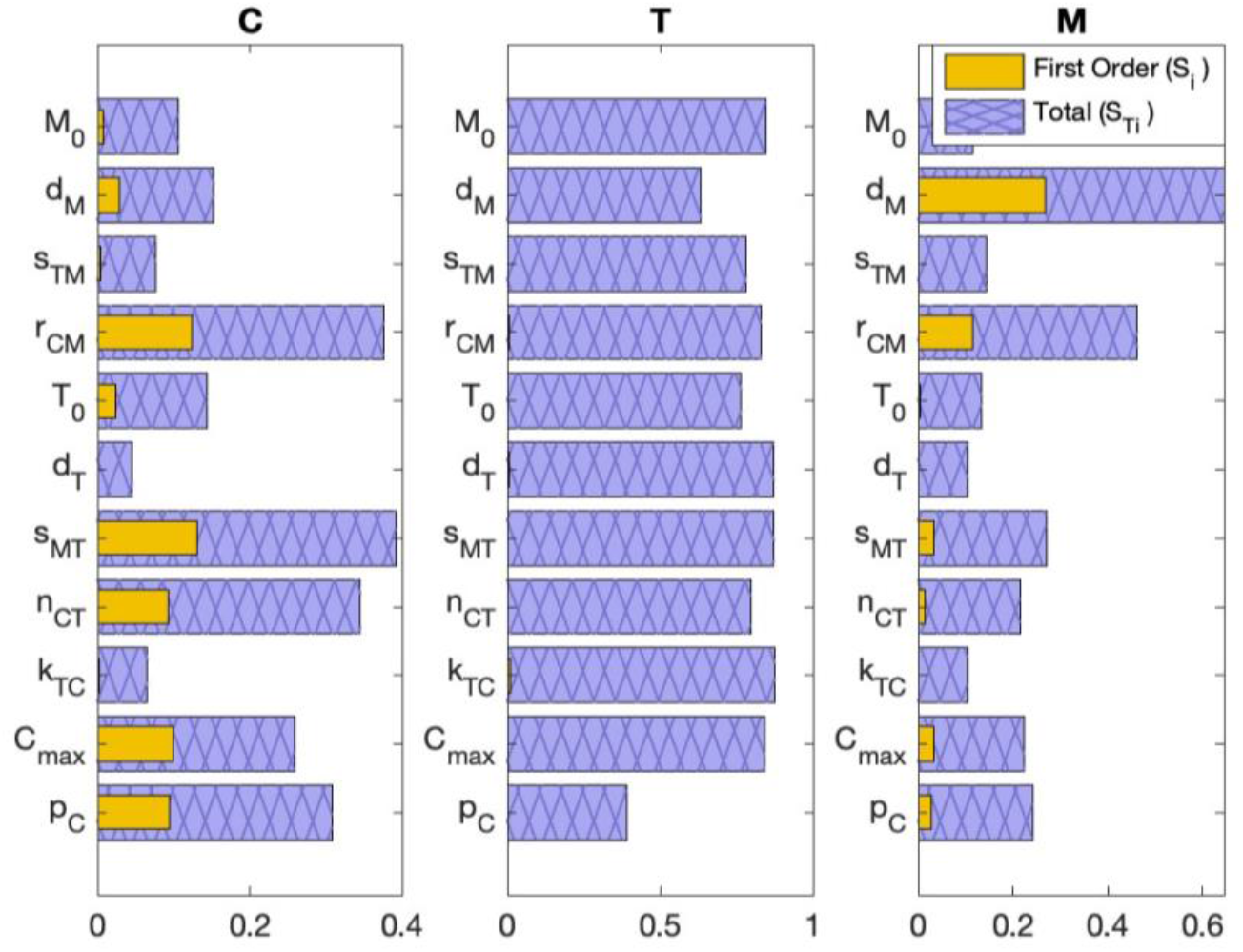
Global sensitivity analysis. eFAST was conducted at day 20 after tumor implantation. Since Fig 4 suggested that most of the tumor is cancer (*C*) rather than T cells (*T*) or MDSCs (*M*), we focused on cancer sensitivity to conclude that practically identifiable **Set 4** (*n*_*CT*_, *r*_*CM*_, and *k*_*GM*_) causes more variation in virtual cohort generation than the other practically identifiable subsets (**Set 1**: *p*_*C*_, *T*_0_, and *k*_*GM*_; **Set 2**: *p*_*C*_, *r*_*CM*_, and *k*_*GM*_; **Set 3**: *k*_*TC*_, *r*_*CM*_, and *k*_*GM*_; **Set 5**: *r*_*CM*_, *d*_*M*_, and *k*_*GM*_) from Section 2.5. This conclusion was based on both the first-order index, *S*_*i*_, and the total order index, *S*_*Ti*_.

Fig 7 shows our results from the eFAST method. The eFAST method produces two measures of sensitivity: the first-order index, *S*_*i*_, and the total order index, *S*_*Ti*_. Here *S*_*i*_ represents the effect that varying a single parameter, *p*_*i*_, has on the model output, while *S*_*Ti*_ = (*D* − *D*_(−*i*)_)/*D*, where *D* is the variance of the model and *D*_(−*i*)_ is the variance of the complementary set (i.e., all parameters other than *p*_*i*_). In other words, *S*_*Ti*_ can be considered the percentage (out of 1) of the model’s variance that is attributed to *p*_*i*_ and *p*_*i*_’s interactions with other parameters that increase the model’s variance (thus, *S*_*Ti*_ ≥ *S*_*i*_). Based on the five sets from Section 2.5 (**Set 1**: *p*_*C*_, *T*_0_, and *k*_*GM*_; **Set 2**: *p*_*C*_, *r*_*CM*_, and *k*_*GM*_; **Set 3**: *k*_*TC*_, *r*_*CM*_, and *k*_*GM*_; **Set 4**: *n*_*CT*_, *r*_*CM*_, and *k*_*GM*_; **Set 5**: *r*_*CM*_, *d*_*M*_, and *k*_*GM*_), we compared the sensitivity of the following parameters: the tumor growth rate (*p*_*C*_), homeostatic T cell population (*T*_0_), MDSC recruitment rate (*r*_*CM*_), T cell kill rate of cancer cells (*k*_*TC*_), cancer-mediated proliferation rate of T cells (*n*_*CT*_), and MDSC death rate (*d*_*M*_).

Across all parameters, T cells show similar sensitivity in terms of *S*_*i*_ and *S*_*Ti*_. Both cancer cells and MDSCs show lower sensitivity to *k*_*TC*_ and *T*_0_ than to the other parameters under consideration, thus eliminating **Sets 1** and **3**. MDSCs show the highest sensitivity to *d*_*M*_ out of all parameters, while cancer cells show lower sensitivity to *d*_*M*_ than to *p*_*C*_, *r*_*CM*_, and *n*_*CT*_. Since Fig 4 suggested that the tumor contains more cancer cells than MDSCs, the sensitivity of the cancer cells was more important in determining which of the five parameter sets is the most sensitive for the total tumor. Thus, we eliminated **Set 5** from consideration. Although we specifically considered sensitivity at day 20, eliminating **Sets 1, 3**, and **5** from consideration was consistent across days 10, 15, and 25 (Supplementary Fig S1). Since the data collection period lasts until day 20, this was a sufficient time range to consider.

The two remaining parameter sets were **Set 2** (*p*_*C*_, *r*_*CM*_, and *k*_*GM*_) and **Set 4** (*n*_*CT*_, *r*_*CM*_, and *k*_*GM*_), where the determining factor was cancer cell sensitivity to the tumor growth rate (*p*_*C*_) and the cancer-mediated proliferation rate of T cells (*n*_*CT*_). In terms of both *S*_*i*_ and *S*_*Ti*_, cancer cells are more sensitive to *n*_*CT*_ than *p*_*C*_ on day 20, 25, and 40, similarly sensitive in terms of *S*_*i*_ at day 15, and are more sensitive to *p*_*C*_ than *n*_*CT*_ in terms of *S*_*i*_ and *S*_*Ti*_ at day 10. These results suggest that **Set 2** or **Set 4** could generate the virtual cohort. Since cancer cells exhibit slightly more sensitivity to *n*_*CT*_ than *p*_*C*_ later in the data collection period when data variability is greater, we concluded that **Set 4** (*n*_*CT*_, *r*_*CM*_, and *k*_*GM*_) should be used for virtual cohort generation.

**Set 4** also exhibited variability experimentally. In vitro migration assays using a bladder cancer cell line ^23^ showed that the cancer-mediated recruitment rate of MDSCs (*r*_*CM*_) varied within the range of 0.0880 to 0.1760 *day*^−1^ (Subsection 1.4.1 of Supplementary Section S1), which approximately corresponds to the *r*_*CM*_ range of accepted virtual mice (*r*_*CM*_ values accepted for the virtual cohort in Section 2.7 ranged from 0 to 0.3 even though *r*_*CM*_ was allowed to vary from 0 to 0.5). As for the Gem kill rate of MDSCs (*k*_*GM*_), flow cytometry analysis showed variability in the MDSCs in the bladder tumors of mice treated with the same dose of Gem 4 days post-treatment (^14^, Fig 4B). Thus, there is variability in Gem’s ability to kill MDSCs. Lastly, ^21^ showed that the percentage of proliferating CD8^+^ T cells can vary widely, with bladder cancer patients exhibiting 1.5% to 15.3% of CD8^+^ T cells expressing the Ki67 marker of cell proliferation. Therefore, the cancer-mediated T cell proliferation rate (*n*_*CT*_) also exhibits intersubject variability clinically.

### 2.7 Curation of the Virtual Cohort

From practical identifiability analysis, five sets of parameters were found to be practically identifiable in terms of ultrasound data alone. Sensitivity analysis showed that the model is more sensitive to **Set 4** (*n*_*CT*_, *r*_*CM*_, and *k*_*GM*_), implying that varying this set better reproduces the variation seen in the data. Experimental data showed that these three parameters all vary within the context of bladder cancer ^14,21,23^. Thus, we have a parameter subset that is practically identifiable, sensitive, and exhibits variability experimentally, so it is suitable for virtual cohort generation.

Based on the “accept-or-reject” method from ^6^ described in Section 4.4, we uniformly sampled 500,000 parameter sets by varying *n*_*CT*_, *r*_*CM*_, and *k*_*GM*_ within their ranges and fixed the other parameters to their best fit values from Supplementary Table S1. Based on the day 6 ultrasound mean (27.6554 *mm*^3^) and standard deviation (11.2253 *mm*^3^), we normally sampled the initial volume at day 6 and then multiplied it by the cell percentages (Fig 4) to obtain 500,000 different initial conditions for the 500,000 plausible mice. Of these plausible mice, 10,424 were within 2 standard deviations of each ultrasound data point for the untreated, Gem, and OT-1 cohorts, while the remaining Gem+OT-1 combination cohort was left out to be used for cohort-level cross-validation in Section 2.8. Fig 8 showcases our virtual murine cohort of 10,424 virtual mice under the four different treatment regimens, where the virtual cohort under the Gem+OT-1 regimen is also included for comparison to experimental data.

**Fig 8.**
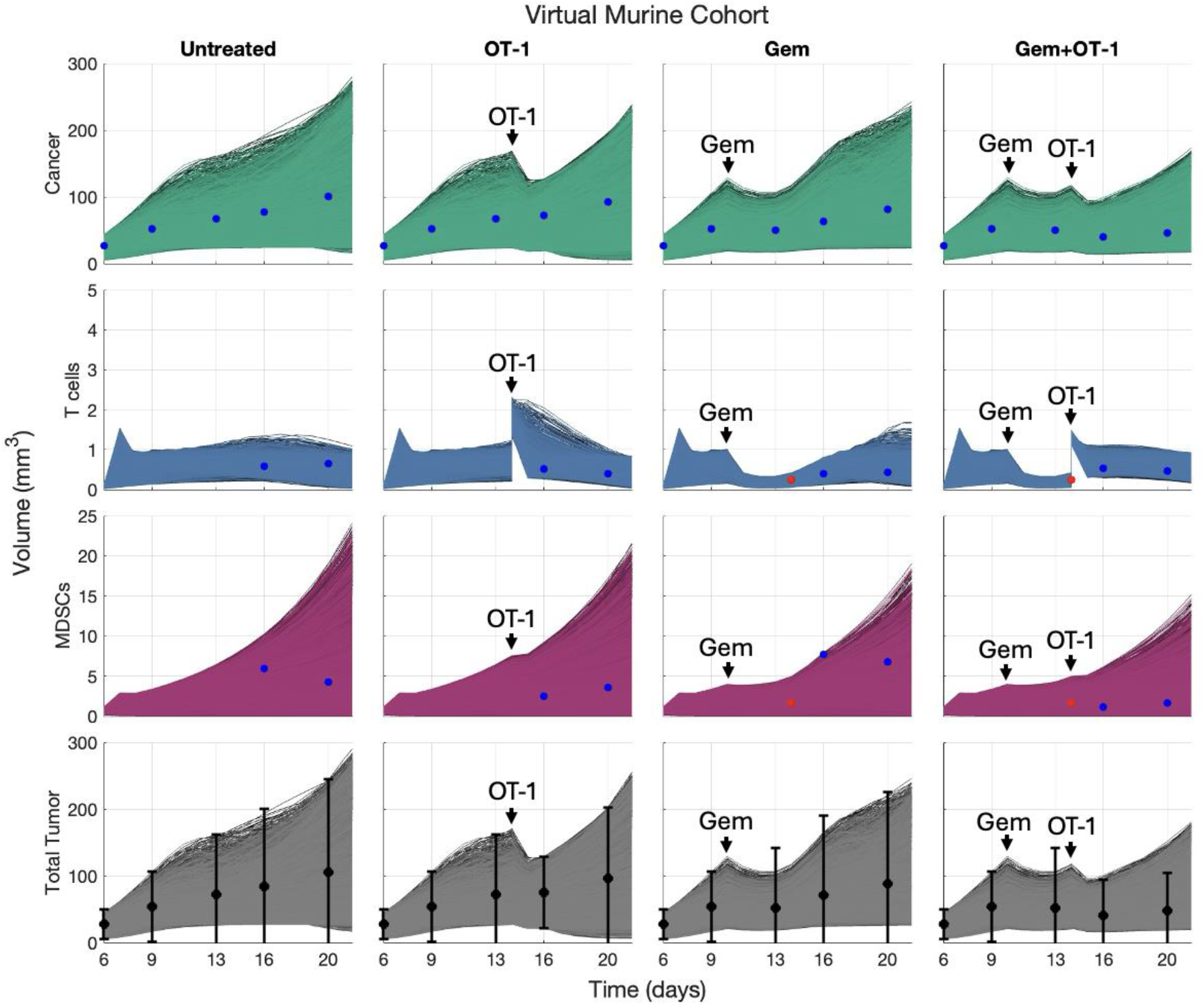
Numerical simulations of virtual cohort under 4 regimens. Virtual cohort of 10,424 mice as determined by the “accept-or-reject” method compared to experimental data. The columns showcase the same 10,424 mice under different treatment regimens. Black dots and error bars represent the mean total tumor size and 2 standard deviations, respectively, from ultrasound data (^14^, Fig 6C). Blue dots represent the histology-modified ultrasound data, while red dots represent ultrasound volumes informed by flow cytometry data.

### 2.8 Validation of Virtual Cohort

The final step is validation of the virtual cohort. In this section, we compared the distributions (Section 2.8.1) and the tumor growth dynamics (Section 2.8.2) of the virtual cohort to the experimental over time.

#### 2.8.1 Cohort Distribution Comparison

We considered “before and after” cohort-level validation and cohort-level cross-validation as was presented in ^1^. In Fig 9, we compared the violin plots of the virtual cohort under therapy versus the experimental cohorts, where the thickness of the violin corresponds to more mice observed at that volume. Similar to ^27^, we used the Kolmogorov-Smirnov test to determine whether distributions from the virtual cohort differed significantly from those of the experimental cohort at the various time points.

**Fig 9.**
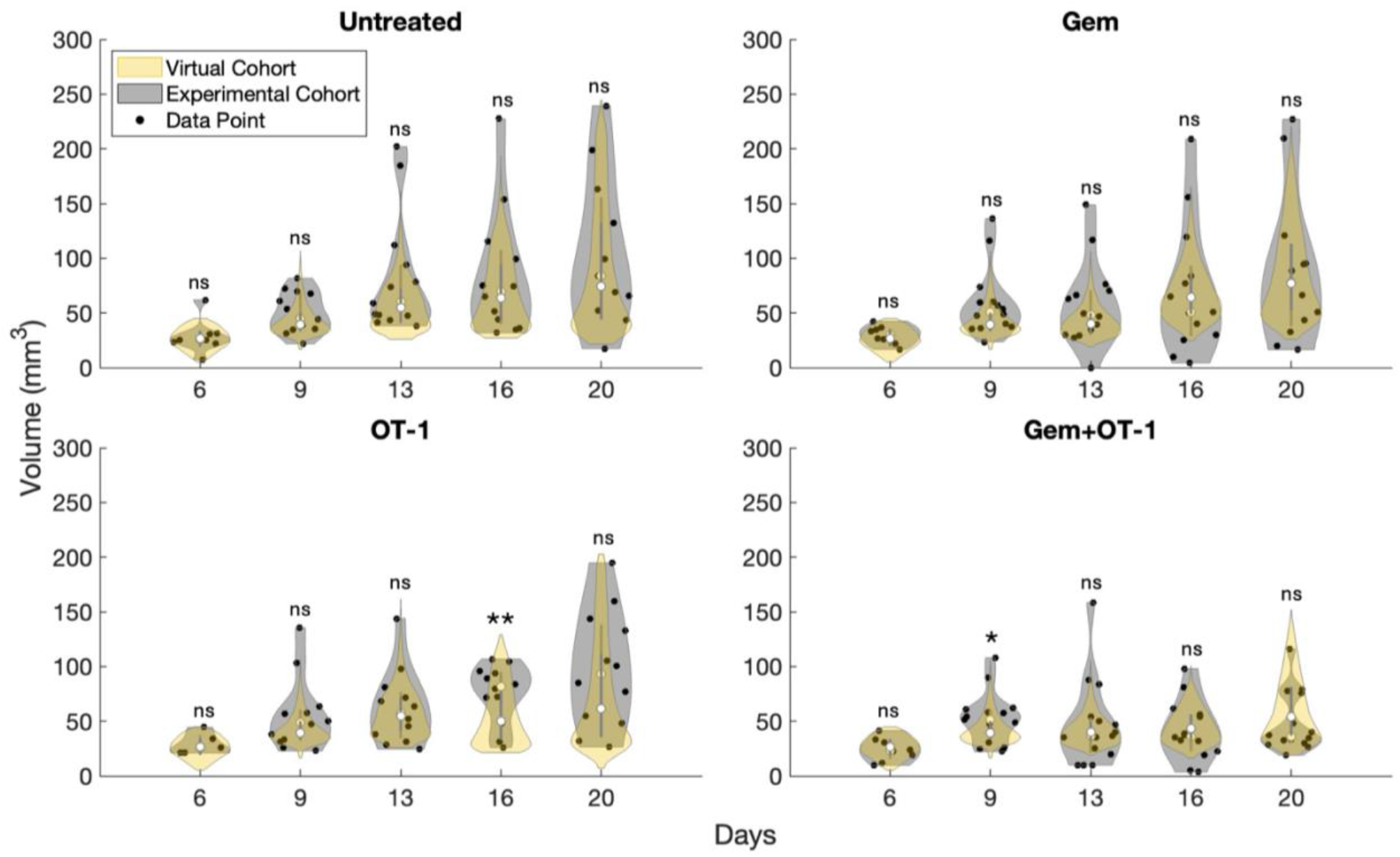
Statistical comparison of virtual cohort to experimental. Violin plots depict the total tumor volume distributions of the virtual and experimental cohorts. White circles along thicker lines inside the violin represent the medians and interquartile ranges, respectively. Results show that only two time points are significantly different between the virtual and experimental cohorts. All statistical analyses were performed using the Kolmogorov-Smirnov test. ns, not significant; *, p ≤ 0.05; **, p ≤ 0.01.

In the “before and after” cohort-level validation, the variability of the virtual cohort is compared to variability of the three experimental cohorts (untreated, Gem monotherapy, and OT-1 monotherapy) that we used to generate the virtual cohort. This comparison was at every time point for which the experimental data was collected. We found that there was no significant difference between the virtual cohort under each therapy and the three experimental cohorts with one exception. For OT-1 monotherapy on day 16, the distributions were significantly different with a p-value of 0.0095. This showed that the virtual cohort initially over predicts the response to OT-1 treatment, which occurs at day 14.

We also considered cohort-level cross-validation by comparing the virtual cohort to the Gem+OT-1 combination cohort to assess the cohorts’ ability to generalize to a cohort not used in its generation. With this data set, there was one significant difference seen at day 9 with a p-value of 0.0229. This result showed that, in general, the virtual cohort can replicate variability seen in a new experimental cohort under these two therapies.

#### 2.8.2 Disease Dynamics Comparison

While the experimental distributions were largely represented by the virtual cohort, we also wanted to ensure that the tumor growth dynamics for each experimental mouse were seen in the virtual cohort. For each experimental mouse, we identified a virtual mouse under the same treatment regimen that best recapitulated the tumor growth dynamics by calculating the least squares error. Results are plotted in Fig 10. Simulations started at day 9 instead of 6 since data collection did not begin for some mice until day 9. Also, for experimental mice that died during treatment, its digital twin’s simulation stopped at the last data collection time point. Fig 10 shows that each mouse has a realistic digital twin within the virtual cohort.

**Fig 10.**
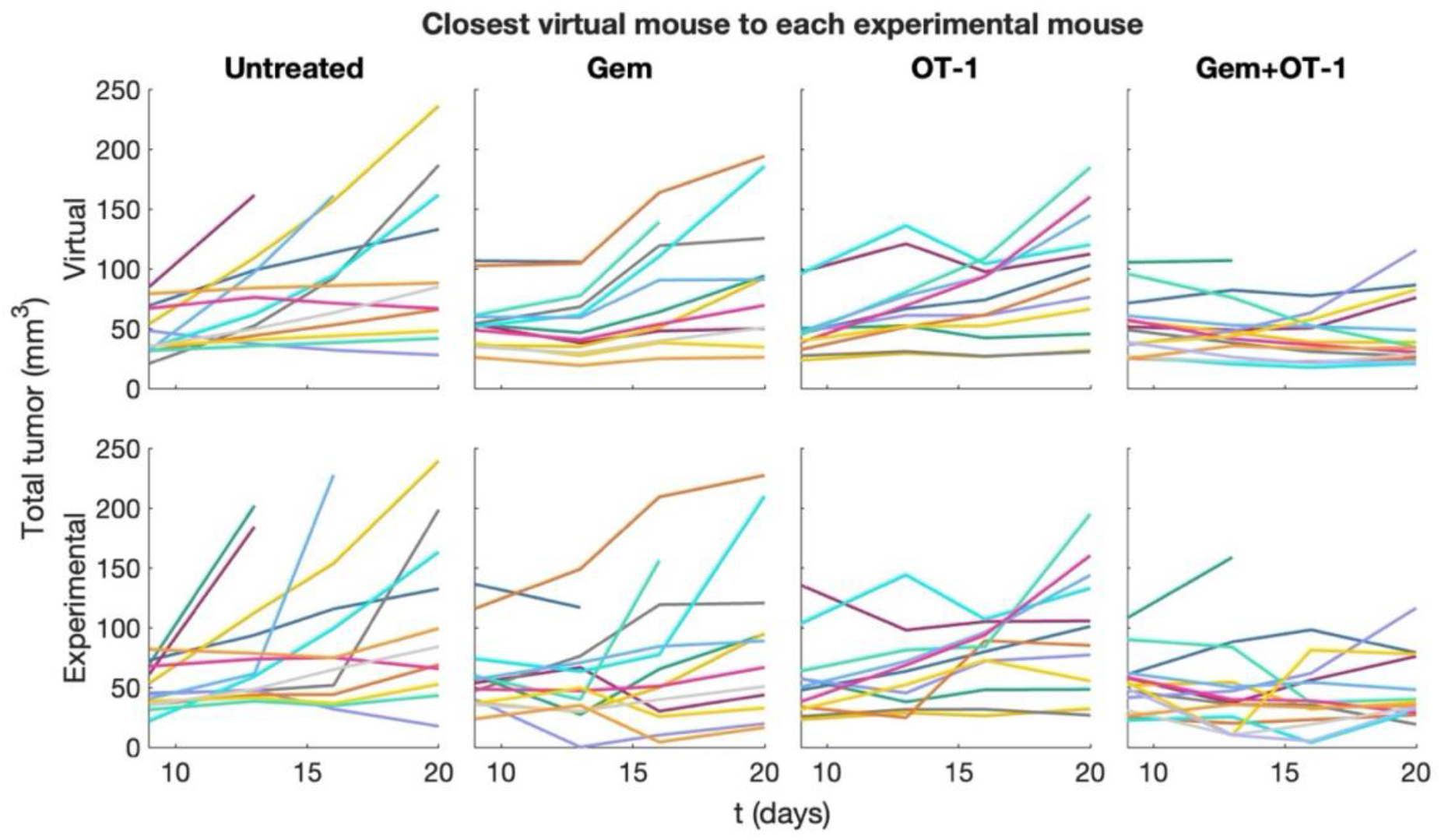
Tumor growth dynamics: virtual cohort versus experimental. For each experimental mouse, we found a realistic digital twin within the virtual cohort that minimizes the least squares error compared to the other virtual mice. The experimental mouse (bottom row) and its corresponding digital twin (top row) are plotted in the same color.

### 2.9 Summary and Guide for Utilizing the Virtual Cohort Pipeline for Other Models

While we used this pipeline for combination therapy in a murine bladder cancer model, this approach can be adapted to other treatments, different diseases, and patient data if the mathematical model implemented is an ordinary differential equations (ODE) model. When developing the fit-for-purpose model, some questions to ask are:

- What are the drivers of disease progression in this disease context?
- Who are the most important cellular/treatment players to include as variables?
- Which mechanisms are essential to accurately depict how the therapy works?
- What data is currently available? What type of additional data can one procure?
- Can this model output an experimentally (or clinically) relevant measurement that can be captured longitudinally with data?

A model that can answer these questions will be biologically and pharmacologically relevant, while its complexity is constrained by available (or possibly available) data. Further, if the model output matches longitudinal data, this opens the possibility of future digital twins and patient (or animal) stratification.

After model development, our next step is a priori structural identifiability analysis, which differs from other virtual cohort pipelines^1–7^ but ensures that the type of data used for parameter estimation can identify parameters given the model’s structure. The differential algebra approach ^25,28^ (or other a priori methods ^29,30^) checks whether the model is *globally* structurally identifiable, meaning that all parameters can, in principle, be *uniquely* determined from the specified data type. If it is not, then either the model’s structure or the type of data used for parameter fitting needs to change. This is where the answer to the question “what type of additional data can one procure?” becomes relevant.

If the model is structurally identifiable given additional data, then the next step is parameter estimation. However, if additional data cannot be retrieved, the model needs to be simplified to work with the available dataset. This may be a complicated process that requires eliminating entire variables, or it could be simple, such as merely combining parameters. Since the differential algebra approach proofs may take some time and require symbolic manipulation software such as Wolfram Mathematica ^31^, future improvements could increase efficiency by using computational tools, such as DAISY ^32^, among others ^33^. Some tools may also assist in reparametrizing structurally unidentifiable models ^34^.

Using the necessary data determined from structural identifiability, one can now estimate parameters. If there are many parameters, those for which one is more confident from the biological literature can be fixed. Parameter bounds in fitting may also ensure that the model is making biologically relevant assumptions.

The next goal is to determine the parameter subset that will generate the virtual cohort. Other pipelines suggest using practical identifiability or sensitivity analysis to identify this subset. Using only one of these options is viable, as we discuss below. However, if the modeler wants a subset that can capture the variability of the experimental data, be utilized to make digital twins in the future, and may be used to stratify digital twins into different treatment groups, the generating parameter set should be both practically identifiable and sensitive. We suggest that practical identifiability analysis precede sensitivity analysis. In this way, one first identifies several parameter subsets that are practically identifiable from longitudinal data and then uses sensitivity analysis to select the subset that maximizes the variability of the virtual cohort.

Practical identifiability analysis determines parameters that can be identified given an experimental data set. If only this analysis is used to find the generating subset, this is still a suitable method since a model will at least be somewhat sensitive to a practically identifiable parameter subset. This result follows from considering the shape of a practically identifiable parameter’s profile likelihood; varying this parameter alters the model’s error relative to the data and, consequently, its output. Thus, these parameters cause at least some variability needed for the virtual cohort, although it may be less variability than our method. Regardless, practical identifiability should be assessed using longitudinal data that can be tracked over time for an individual, as this data is likely used in virtual cohort sampling, so these parameters are suitable for comparison with that data set. Further, if longitudinal data can be used to identify the parameter subset, this enables the future generation of digital twins.

If there are issues finding a practically identifiable subset from profile likelihoods, one should consider reducing or changing the parameter subset tested, as interactions between parameters may be causing a lack of identifiability. The rank of the Fisher Information Matrix (FIM) can be used to gauge how many parameters should be practically identifiable, and ^35^ presents a method for using the FIM to identify parameter subsets that may be identifiable. However, there are mixed reviews on the FIM— particularly for models that are more complicated than a linear regression model ^13,28^, so practical identifiability should be checked with profile likelihoods; this follows the method in ^35^. Alternatively, if practical identifiability and sensitivity analysis are both used to determine the virtual cohort’s generating parameter subset, one could perform them in parallel and test sensitive parameters, as these are more likely to be practically identifiable. In this case, sensitivity analysis would still ultimately follow practical identifiability analysis to determine which discovered practically identifiable subset causes the most variation in the model output.

Otherwise, if only sensitivity analysis is used to determine the generating parameter subset, this may result in a larger number of parameters being used to generate the virtual cohort, thereby increasing variability in individual treatment responses. In general, this is a good avenue to pursue. However, there may be an issue if treatment optimization indicates that the virtual cohort should be stratified into subgroups based on improved response to alternative scheduling or dosing strategies. While virtual patients can be stratified into treatment subgroups, it is difficult to determine the treatment subgroup an actual patient should be in, since the parameters used to fit the patient’s data are not practically identifiable and thus may not be unique. Also, when using a larger generating parameter subset, as the number of parameters increases, the acceptance rate for plausible patients decreases, a phenomenon known as the curse-of-dimensionality ^36^. The lower acceptance rate makes it more difficult to generate a virtual cohort, especially if each plausible patient is compared to several experimental cohorts. Lastly, choosing the threshold that distinguishes sensitive from insensitive parameters is somewhat arbitrary.

As for the sensitivity analysis method, we used eFAST, which is a variance-based global sensitivity analysis (similar to the Sobol method ^37^), but eFAST requires fewer model evaluations than the Sobol method thus improving upon the run time ^38,39^. Unlike the partial rank correlation coefficient (PRCC) method, eFAST does not require monotonicity between parameter input and model output, so this also makes eFAST a method that can work regardless of your model as that condition does not need to be verified ^40^. So, in terms of both speed and convenience, eFAST is a good option to choose, although other sensitivity analysis methods could be considered, such as Morris or LH-OAT ^39^.

After obtaining the generating parameter subset and confirming its ability to reflect biological or clinical variability, these parameters are randomly sampled within their respective ranges, and parameter sets that fall within a specified threshold from each data point become the virtual cohort. This is referred to as the “accept-or-reject” method ^6^. Determining a threshold for acceptance goes hand-in-hand with the “before and after” cohort validation step, which compares the distributions of the virtual cohort with those of the experimental cohorts at each data collection time point. Some potential error thresholds to try are 2, 3, or 4 standard deviations from the mean or the range from the minimum to maximum data point at each time. While a larger error threshold captures more variability, if the distributions of the virtual and experimental cohorts are no longer similar, it may be more important to reflect the distributions of the data than to over-represent outliers. Future improvements to our pipeline could include a prevalence-weighting method ^41^ to further improve the ability of the virtual cohort to recapitulate the data’s distributional shape over time, not just its range.

Lastly, “before and after” validation using the Kolmogorov-Smirnov test can confirm that the distribution of the virtual cohort is not significantly different from that of the experimental cohort. It is essential that these distributions are similar. If the virtual cohort is merely within the bounds of the experimental data, certain subgroups in the data could be over-represented. This may become an issue when treatment regimens are tested on the virtual cohort, since a regimen may prove more robust there, but in reality, the virtual cohort has a skewed representation, and thus, the regimen may not be robust in an experimental cohort. To test how the virtual cohort might perform under different treatment regimens, if there are enough experimental cohorts in the data set, we advise modelers to leave one out of the “accept-or-reject” step and statistically compare that cohort’s data to the virtual cohort under the same regimen. This second check is called cohort-level cross-validation. A third and final validation can focus on the individual level to ensure that the disease is progressing realistically by confirming that the dynamics over time observed in the experimental cohort are also observed in the virtual cohort.

By using our virtual cohort pipeline, a modeler can better capture the dynamics of each variable (a priori structural identifiability), produce digital twins using longitudinal data (practical identifiability), capture interpatient (or intersubject) variability (sensitivity analysis), and validate that the variability seen in the virtual cohort matches that of the experimental data. In the future, a modeler’s virtual cohort can be used to optimize and evaluate treatment robustness or to stratify individuals by regimen using digital twins. The simple act of improving the scheduling and dosing of current therapies using virtual cohorts can help more patients respond better to therapies that are already available. Further, these virtual cohorts help expand small data sets in early-stage clinical trials or preclinical experiments, thereby maximizing the efficacy of therapies early on, given patient (or animal) variability.

## 3 Discussion

In this paper, we took an experimental data set of 55 mice and generated a virtual cohort consisting of over 10,000 virtual mice to capture the variability in OT-1 and Gem treatment using the pipeline detailed in Fig 1. Validation using the Kolmogorov-Smirnov test compared the variability of the virtual cohort with that of the four experimental cohorts, showing that the test could not distinguish between the virtual and experimental cohorts at 18 of 20 time points (Section 2.8). Our virtual cohort also more realistically captured T cells and MDSCs in the tumor microenvironment over time, as structural identifiability necessitated data on individual cell types for the initial parameter fitting (Section 2.3–2.4). This guided us to use supplemental histology and flow cytometry to determine the percentages of T cells and MDSCs in the tumor microenvironment over time, and then to use this information to modify ultrasound data. Since both histology and flow cytometry data require euthanizing the mouse to retrieve the data, we wanted future iterations of the model to require only data that can be measured longitudinally with ultrasound (Section 2.5). Using profile likelihoods, we determined subsets of parameters that can be identified using ultrasound data alone; thus, these subsets open the opportunity to predict treatment response in mice in real time in the case that optimized regimens for the virtual cohort are tested experimentally.

After identifying five practically identifiable parameter subsets, our next question was: *which subset causes the most variability in the model output?* The subset that answered this question would be best at capturing the intersubject variability of the experimental data. Global sensitivity analysis with eFAST showed that the tumor was most sensitive to the fourth subset: cancer-mediated T cell proliferation rate (*n*_*CT*_), cancer-mediated MDSC recruitment rate (*r*_*CM*_), and the Gem kill rate of MDSCs (*k*_*GM*_) (Section 2.6). Further, experimental data showed variability in these parameters ^14,21,23^. Thus, since this subset (1) was suitable to compare to ultrasound data for our virtual cohort sampling, (2) could be used for the future creation of digital twins (i.e., it is practically identifiable with longitudinal data), (3) caused variability mathematically as seen from sensitivity analysis, and (4) showed variability experimentally, it was suitable to generate the virtual cohort.

Varying these three parameters within their ranges reported in the literature, we used the “accept or reject” method ^6^ to accept virtual mice into the virtual cohort (Section 2.7). Our acceptance threshold was 2 standard deviations from the data mean at each time point for the untreated, Gem monotherapy, and OT-1 monotherapy experimental cohorts, as this threshold better reproduced the experimental summary statistics than thresholds of 1, 3, or 4 standard deviations. 10,424 of the 500,000 tested mice produced simulations within 2 standard deviations at each data point across the three experimental cohorts and were accepted into our virtual cohort.

Validation occurred in three steps (Section 2.8). The first step was “before and after” validation, in which we compared the virtual cohort with the three experimental cohorts used to generate it. In this step, the Kolmogorov-Smirnov test identified only one significant difference out of 15 time points between the distributions of the virtual cohort and the three experimental cohorts. The second validation was a cohort-level cross-validation, where we compared the virtual cohort to an experimental cohort not used in its generation, namely the Gem+OT-1 combination cohort. Results from this second validation showed that the Kolmogorov-Smirnov test was unable to distinguish whether the virtual and experimental cohorts were from different populations at all but one of the data points. Although cross-validation was performed on only one experimental cohort, this initial test suggested that the virtual cohort could be used under a different regimen with about 80% accuracy. We also verified that the tumor progression dynamics of the virtual cohort were similar to those of the experimental cohort by identifying a digital twin within the virtual cohort that best reproduced the data for each experimental mouse and then plotting the simulation dynamics against the data.

Future improvements to the virtual cohort pipeline would include prevalence weighting in virtual cohort sampling to ensure that the variation observed in the experimental cohorts is even more closely replicated by the virtual cohort. Also, using a numerical form of a priori structural identifiability analysis could help speed up the virtual cohort pipeline by removing the need for additional theorems and proofs.

In the future, alternative treatment regimens can be tested to find the regimen that works best for most individuals in the virtual cohort. This could be done in several ways, including trial and error, optimizing for the parameter set representing an average individual and testing that regimen on the virtual cohort, or optimizing across the entire virtual cohort. Alternatively, we could fit the practically identifiable parameter subset to the longitudinal ultrasound data for each experimental mouse, thereby creating a digital twin. Then, we could optimize treatment for each individual and test the optimal regimens on the virtual cohort to determine the most robust regimen. Doing it this way would ensure that the regimen eventually tested experimentally is theoretically optimal for an individual who has existed. This further guards against any unfair weighting that could result from optimizing the regimen for the virtual cohort itself. When testing these treatment schedules, several regimens may emerge as the best for different subgroups of the virtual cohort, necessitating the need to stratify experimental mice for better outcomes. When the regimen is tested experimentally, the practically identifiable parameter set (*n*_*CT*_, *r*_*CM*_, *k*_*GM*_) can be fit to an individual’s longitudinal ultrasound data, thereby creating a digital twin. These digital twins can be used to personalize predictions of disease progression and stratify mice into different treatment subgroups to improve response.

While we applied our virtual cohort pipeline to preclinical mice data, a similar approach can be used to design a virtual patient cohort. The orthotopic MB49-OVA bladder tumor in mice is considered a preclinical model of non-muscle invasive bladder cancer (NMIBC). Moreover, the experimental procedures of adoptive therapy with OT-1 T cells mimic the clinical trial application of intravesical adoptive cell therapy with tumor-infiltrating lymphocytes (TIL) in NMIBC patients (NCT05768347). Since most NMIBC patients undergo transurethral resection of bladder tumor (TURBT) at the start of treatment ^42^, with multiple follow-up procedures and treatments, a possible research plan would be to collect longitudinal patient data and organize it according to the patient’s time since TURBT. Common surveillance data collected during patient visits include a urine cytology from a urine sample, cross-sectional imaging, and a cystoscopy, where a clinician uses a thin camera scope to evaluate the lining of the bladder and urethra ^42^. This data is taken longitudinally for patients according to the AUA guidelines ^42^.

This data could be used to develop a virtual NMIBC patient cohort. Clinicians routinely give a size approximation for any suspect tumor in a cystoscopy and take a biopsy if needed, which could be analyzed for TIL or other immune cells. Since this data may be sparse if no tumor is found on cystoscopy, virtual cohort generation may require additional data elements. Next generation urine analysis of utDNA (urinary tumor DNA) ^43^, FDA-approved protein-based biomarkers for bladder cancer (such as NMP22) ^44^, immune cells, or cytokines could be used to add further refinement to the model. Using this data may require the bladder tumor-immune model be altered to include a urine compartment. With a virtual NMIBC patient cohort developed like so, various treatment regimens could be tested directly on virtual patients instead of extrapolating treatment robustness results from murine experiments. Thus, the data variability captured by the virtual cohort pipeline can enable the identification of treatment schedules that yield better outcomes for more patients.

Our work shows that virtual cohorts model the variability in treatment response and disease progression observed in experimental or clinical data while generating larger cohorts. These cohorts may help design more robust treatment protocols that better suit more individuals, even in the context of the limited data sets of preclinical studies or early-phase clinical trials. Further, patient stratification using identifiable parameter sets can place an individual in a treatment cohort that may allow them to receive better therapy for their specific situation. These features can support researchers with limited data and guide them toward the next steps in advancing drug development and patient care.

## 4 Methods

### 4.1 Differential algebra approach

The differential algebra approach is an a priori, global structural identifiability method for nonlinear ordinary differential equation models ^25^. Given the structure of a model, a priori structural identifiability analysis aims to determine if a certain type of data can determine model parameters. By definition, a parameter, *p*_*i*_, is globally structurally identifiability if, for an observed output, there is a unique value, 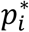, that results in that output. A model is globally structurally identifiable if all parameters are globally structurally identifiable.

A simple example of the differential algebra approach can be found in ^28^. The steps involved are as follows: first, one determines which variable(s) have existing data. Then, the system is rearranged to be only in terms of these variable(s) and their derivatives. These new polynomials are known as the “input-output relation(s)” or the “characteristic set.” The coefficients of the input-output relation(s) form a set of identifiable combinations of parameters, which can be used to solve for each parameter. For instance, if *a* and *b* are parameters from the model and *a* + *b* and *a* − *b* are identifiable combinations, *a* can be solved for:

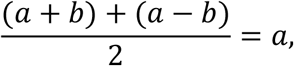

and thus *b* can be solved for. The parameters that can be solved analytically are globally structurally identifiable. If this is true of all parameters, then the model is globally structurally identifiable, and consequently, the existing data type is suitable for parameter fitting.

### 4.2 Profile likelihood method

Profile likelihoods maximize the likelihood of a parameter, *p*, by fixing *p* to different values within its range and then profiling the remaining parameters to maximize the likelihood function. Assuming that the errors between the model predictions and the experimental data are normally distributed with a mean of 0 (which assumes that, on average, model predictions are accurate), one can conclude that maximizing the likelihood is equivalent to minimizing the least squares error (LSE) ^45^.

To produce profile likelihoods, one first chooses which model parameters to profile: *p*_1_, …, *p*_*n*_. Using these parameters with other model parameters fixed, the best fit, 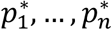, compared to the data is determined in terms of LSE. Then, each parameter, 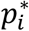 is varied within a specified range of its best fit value, *p*^∗^, each time refitting the other parameters *p*_1_, …, *p*_*i*−1_, *p*_*i*+1_ …, *p*_*n*_ to the data set and calculating each new fit’s LSE. The plot of the LSE of these fits with respect to the value of *p*_*i*_ is *p*_*i*_’s profile likelihood. If this plot exhibits a distinct minimum, this indicates that there is a unique parameter value that best represents the data, thus that parameter is practically identifiable. The threshold for the 95% confidence interval is also plotted, which is determined using the chi-squared distribution, *χ*, at a confidence level of *α* = 0.05:

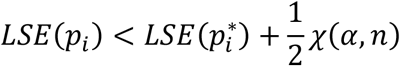

where *n* represents the degrees of freedom corresponding to the *n* chosen parameters. The intersection of the profile likelihood with this error threshold is the 95% confidence interval, which allows one to be 95% confident that the true parameter value falls within this interval. A smaller 95% confidence interval indicates higher confidence in the parameter estimate.

### 4.3 eFAST method

If a variable is sensitive to a parameter, this signifies that changing that parameter’s value causes fluctuations in the variable output. We performed global sensitivity analysis using the extended Fourier Amplitude Sensitivity Test (eFAST), which was originally proposed by ^46^ and further developed by ^38^ to include a calculation of total order sensitivity indices (*S*_*Ti*_), which includes the effect that interactions between parameters has on the model’s sensitivity. MATLAB code for eFAST was developed by the Kirshner Lab at the Univeristy of Michigan and is publically available ^47^.

To perform eFAST, one first samples parameters uniformly from the unit hypercube using the uniform search curve ^38^. Based on the number of model parameters, ^38^ [Tab. 2, Fig 4] gives suggestions for the number of resamplings and the number of parameter combinations per resampling to ensure that the space is thoroughly and evenly sampled. For instance, we performed 2 resamplings each containing 1,977 combinations of parameter values, which was the sampling size suggested for 11 parameters to produce accurate estimates compared to the sensitivity indices analytical values. After sampling, parameter values are scaled from the unit hypercube to their ranges listed in Table S1, and the variance is determined using model simulations for each parameter set.

### 4.4 “Accept-or-reject” method

The “accept-or-reject” method ^6^ is a variation of the Approximate Bayesian Computation (ABC) rejection method ^36^. Parameters are first sampled within their specified ranges— often a uniform sampling unless a specified prior distribution for the parameter is known. Then, the model is simulated for each parameter set, and the model output error compared to the experimental data is calculated. Parameter sets that produce an error within a specified threshold are then accepted, while the remaining are rejected. Commonly, the error threshold for the ABC rejection method is an arbitrary user-defined value aimed at accepting a small percentage of simulations in order to generate posterior distributions for parameters ^48–50^. However, in “accept-or-reject” method ^6^, one uses a data-informed error threshold for each data point by considering the variation in the data itself at each time point. In ^6^, the authors accepted parameter sets within 3 standard deviations of the data mean at each time point. In constrast, we set the absolute error threshold to be 2 standard deviations at each ultrasound data point as this allowed the virtual cohort to better capture the experimental summary statistics in Section 2.8. These accepted parameter sets then become the virtual subjects.

## Acknowledgements

This work was supported by the US National Institutes of Health, National Cancer Institute grant R01-CA259387 (to KR and SPT) as well as the Department of Defense grants W81XWH-22-1-0339, W81XWH-22-1-0340, and W81XWH-22-1-0341 (to MP, KR, and SPT). This work was also supported in part by the Shared Resources at the H. Lee Moffitt Cancer Center & Research Institute an NCI designated Comprehensive Cancer Center under the grant P30-CA076292 from the National Institutes of Health. The funders played no role in study design, data collection, analysis and interpretation of data, or the writing of this manuscript.

## Data availability

The data and code generated for this study is available from the GitHub repositories: github.com/HannahGrace314/Virtual-Cohort-pipeline-bladder-cancer and github.com/rejniaklab/Virtual-Cohort-pipeline-bladder-cancer

## Author contributions

HGA and KAR conceptualized the model. HGA developed the pipeline, implemented the methodology, coded the software and figure visualization, and prepared the original draft. SB curated the experimental data. DJN and MAP provided clinical feedback for extending the virtual cohort pipeline to an NMIBC patient population. KAR and SPT assisted in funding acquisition and supervision. All authors read and approved the final manuscript.

## References

1 Chase, J. G. et al. Next-generation, personalised, model-based critical care medicine: a state-of-the art review of in silico virtual patient models, methods, and cohorts, and how to validation them. Biomed Eng Online 17, 24 (2018). 10.1186/s12938-018-0455-y

2 Viceconti, M. et al. Credibility of in silico trial technologies—a theoretical framing. IEEE journal of biomedical and health informatics 24, 4–13 (2019).

3 Sinisi, S., Alimguzhin, V., Mancini, T., Tronci, E. & Leeners, B. Complete populations of virtual patients for in silico clinical trials. Bioinformatics 36, 5465–5472 (2020).

4 Arsène, S. et al. in High performance computing for drug discovery and biomedicine 51–99 (Springer, 2023).

5 Craig, M., Gevertz, J. L., Kareva, I. & Wilkie, K. P. A practical guide for the generation of model-based virtual clinical trials. Front Syst Biol 3, 1174647 (2023). 10.3389/fsysb.2023.1174647

6 Gevertz, J. L. & Wares, J. R. Assessing the Role of Patient Generation Techniques in Virtual Clinical Trial Outcomes. Bull Math Biol 86, 119 (2024). 10.1007/s11538-024-01345-6

7 Kleeberger, J. A. Virtual Patients in Clinical Trials for Drug Development: A Narrative Review. Cureus 17, e85380 (2025). 10.7759/cureus.85380

8 Lu, E.-H., Rusyn, I. & Chiu, W. A. Incorporating new approach methods (NAMs) data in dose–response assessments: The future is now! Journal of Toxicology and Environmental Health, Part B 28, 28–62 (2025).

9 Mirlohi, M. S., Yousefi, T., Aref, A. R. & Seyfoori, A. Integrating New Approach Methodologies (NAMs) into Preclinical Regulatory Evaluation of Oncology Drugs. Biomimetics 10, 796 (2025).

10 Kovatchev, B. P., Breton, M., Dalla Man, C. & Cobelli, C. (SAGE Publications Sage CA: Los Angeles, CA, 2009).

11 ASME. V&V 40 - 2018: Assessing Credibility of Computational Modeling through Verification and Validation: Application to Medical Devices. (2018).

12 Allen, R., Rieger, T. R. & Musante, C. J. Efficient generation and selection of virtual populations in quantitative systems pharmacology models. CPT: pharmacometrics & systems pharmacology 5, 140–146 (2016).

13 Wieland, F.-G., Hauber, A. L., Rosenblatt, M., Tönsing, C. & Timmer, J. On structural and practical identifiability. Curr Opin Syst Biol 25, 60–69 (2021). 10.1016/j.coisb.2021.03.005

14 Bazargan, S. et al. Targeting myeloid-derived suppressor cells with gemcitabine to enhance efficacy of adoptive cell therapy in bladder cancer. Front Immunol 14, 1275375 (2023). 10.3389/fimmu.2023.1275375

15 Yang, Y., Li, C., Liu, T., Dai, X. & Bazhin, A. Myeloid-derived suppressor cells in tumors: from mechanisms to antigen specificity and microenvironmental regulation. Front Immunol 11 (2020). 10.3389/fimmu.2020.01371

16 Clarke, S. R. et al. Characterization of the ovalbumin-specific TCR transgenic line OT-I: MHC elements for positive and negative selection. Immunol Cell Biol 78, 110–117 (2000). 10.1046/j.1440-1711.2000.00889.x

17 Robbins, P. F. Tumor-Infiltrating Lymphocyte Therapy and Neoantigens. Cancer J 23, 138–143 (2017). 10.1097/PPO.0000000000000267

18 Mackey, J. R. et al. Gemcitabine transport in xenopus oocytes expressing recombinant plasma membrane mammalian nucleoside transporters. J Natl Cancer Inst 91, 1876–1881 (1999). 10.1093/jnci/91.21.1876

19 Klysz, D. D. et al. Inosine induces stemness features in CAR-T cells and enhances potency. Cancer Cell 42, 266–282 e268 (2024). 10.1016/j.ccell.2024.01.002

20 Matsumura, N. et al. The prognostic significance of human equilibrative nucleoside transporter 1 expression in patients with metastatic bladder cancer treated with gemcitabine-cisplatin-based combination chemotherapy. BJU Int 108, E110–116 (2011). 10.1111/j.1464-410X.2010.09932.x

21 Blessin, N. C. et al. Prevalence of proliferating CD8+ cells in normal lymphatic tissues, inflammation and cancer. Aging (Albany NY) 13, 14590–14603 (2021). 10.18632/aging.203113

22 Srivastava, M. K., Sinha, P., Clements, V. K., Rodriguez, P. & Ostrand-Rosenberg, S. Myeloid-derived suppressor cells inhibit T-cell activation by depleting cystine and cysteine. Cancer Res 70, 68–77 (2010). 10.1158/0008-5472.CAN-09-2587

23 Zhang, H. et al. CXCL2/MIF-CXCR2 signaling promotes the recruitment of myeloid-derived suppressor cells and is correlated with prognosis in bladder cancer. Oncogene 36, 2095–2104 (2017). 10.1038/onc.2016.367

24 Klein, C. et al. Myeloid-Derived suppressor cells in bladder cancer: an emerging target. Cells 13, 1779 (2024). 10.3390/cells13211779

25 Audoly, S., Bellu, G., D’Angio, L., Saccomani, M. P. & Cobelli, C. Global identifiability of nonlinear models of biological systems. IEEE Trans Biomed Eng 48, 55–65 (2001). 10.1109/10.900248

26 Heinrich, M., Rosenblatt, M., Wieland, F.-G., Stigter, H. & Timmer, J. On structural and practical identifiability: Current status and update of results. Curr Opin Syst Biol 41 (2025). 10.1016/j.coisb.2025.100546

27 Kim, E., Rebecca, V. W., Smalley, K. S. & Anderson, A. R. Phase i trials in melanoma: A framework to translate preclinical findings to the clinic. European Journal of Cancer 67, 213–222 (2016).

28 Eisenberg, M. C. & Jain, H. V. A confidence building exercise in data and identifiability: Modeling cancer chemotherapy as a case study. J Theor Biol 431, 63–78 (2017). 10.1016/j.jtbi.2017.07.018

29 Anstett-Collin, F., Denis-Vidal, L. & Millérioux, G. A priori identifiability: An overview on definitions and approaches. Annu Rev Control 50, 139–149 (2020). 10.1016/j.arcontrol.2020.10.006

30 Chis, O.-T., Banga, J. R. & Balsa-Canto, E. Structural identifiability of systems biology models: a critical comparison of methods. PloS one 6, e27755 (2011). 10.1371/journal.pone.0027755

31 Mathematica (Wolfram Research, Inc., Champaign, Illinois, 2025).

32 Bellu, G., Saccomani, M. P., Audoly, S. & D’Angiò, L. DAISY: A new software tool to test global identifiability of biological and physiological systems. Comput Methods Programs Biomed 88, 52–61 (2007). 10.1016/j.cmpb.2007.07.002

33 Rey Barreiro, X. & Villaverde, A. F. Benchmarking tools for a priori identifiability analysis. Bioinform 39, btad065 (2023). 10.1093/bioinformatics/btad065

34 Joubert, D., Stigter, J. D. & Molenaar, J. An efficient procedure to assist in the re-parametrization of structurally unidentifiable models. Math Biosci 323, 108328 (2020). 10.1016/j.mbs.2020.108328

35 Eisenberg, M. C. & Hayashi, M. A. Determining identifiable parameter combinations using subset profiling. Math Biosci 256, 116–126 (2014). 10.1016/j.mbs.2014.08.008

36 Sunnåker, M. et al. Approximate bayesian computation. PLoS Comput Biol 9, e1002803 (2013). 10.1371/journal.pcbi.1002803

37 Sobol, I. M. Global sensitivity indices for nonlinear mathematical models and their Monte Carlo estimates. Math Comput Simul 55, 271–280 (2001). 10.1016/S0378-4754(00)00270-6

38 Saltelli, A., Tarantola, S. & Chan, K.-S. A quantitative model-independent method for global sensitivity analysis of model output. Technometrics 41, 39–56 (1999). 10.1080/00401706.1999.10485594

39 Wang, A. & Solomatine, D. P. Practical experience and framework for sensitivity analysis of hydrological models: six methods, three models, three criteria. Hydrol Earth Syst Sci Discuss 2018, 1–34 (2018). 10.5194/hess-2018-78

40 Marino, S., Hogue, I. B., Ray, C. J. & Kirschner, D. E. A methodology for performing global uncertainty and sensitivity analysis in systems biology. J Theor Biol 254, 178–196 (2008). 10.1016/j.jtbi.2008.04.011

41 Huang, L. et al. A Proxy-guided Workflow for Virtual Population Development. AAPS J 27, 162 (2025). 10.1208/s12248-025-01134-6

42 Holzbeierlein, J. M. et al. Diagnosis and treatment of non-muscle invasive bladder cancer: AUA/SUO guideline: 2024 amendment. J Urol 211, 533–538 (2024). 10.1097/JU.0000000000003846

43 Linscott, J. A. et al. From detection to cure–Emerging roles for urinary tumor DNA (utDNA) in bladder cancer. Curr Oncol Rep 26, 945–958 (2024). 10.1007/s11912-024-01555-0

44 Yang, Z., Song, F. & Zhong, J. Urinary biomarkers in bladder cancer: FDA-approved tests and emerging tools for diagnosis and surveillance. Cancers (Basel) 17, 3425 (2025). 10.3390/cancers17213425

45 Huang, H.-H. & He, Q. in International Encyclopedia of Education 558–567 (Elsevier, 2023).

46 Cukier, R., Fortuin, C., Shuler, K. E., Petschek, A. & Schaibly, J. H. Study of the sensitivity of coupled reaction systems to uncertainties in rate coefficients. I Theory. J Chem Phys 59, 3873–3878 (1973). 10.1063/1.1680571

47 Kirschner, D. Uncertainty and sensitivity functions and implementation, <http://malthus.micro.med.umich.edu/lab/usadata/ (2008).

48 Ross, R. J. et al. Using approximate Bayesian computation to quantify cell–cell adhesion parameters in a cell migratory process. npj Syst Biol Appl 3 (2017). 10.1038/s41540-017-0010-7

49 Anderson, H. G. et al. Global stability and parameter analysis reinforce therapeutic targets of PD-L1-PD-1 and MDSCs for glioblastoma. J Math Biol 88, 10 (2023). 10.1007/s00285-023-02027-y

50 Lintusaari, J., Gutmann, M. U., Dutta, R., Kaski, S. & Corander, J. Fundamentals and recent developments in approximate Bayesian computation. Syst Biol 66, e66–e82 (2017). 10.1093/sysbio/syw077

