## Supplementary Information 1 for "A virtual cohort framework with applications to adoptive cell therapy in bladder cancer"

**Contents:**

S1. Estimated values for treatment injections and parameters

S1. Table of bladder cancer model parameters for Gem and OT-1 therapy

S1.1. Treatment injections

S1.2. Bladder cancer parameter values

S1.3. T cell parameter values

S1.4. MDSC parameter values

S1.5. Gem Parameter Values

S2. Equilibrium and stability analysis

S3. Global sensitivity analysis figures at alternative times (Fig S1(a-c))

### S1 Estimated values for treatment injections and parameters

The following provides additional information about calculations for treatment injections (S1.1) and parameter values (S1.2-S1.4, listed in Table S1). Some parameter values from literature were converted to be in terms of the units used in our model, while other values used calculations directly from data to ensure the model used realistic estimates.

**Table S1. Bladder cancer model parameters for Gem and OT-1 therapy.**

| LABEL | DESCRIPTION | UNITS | LITERATURE VALUE | CITATION | RANGE FOR ESTIMATION | FINAL SET |
|---|---|---|---|---|---|---|
| **CANCER ($C$)** | | | | | | |
| $\boldsymbol{p_C}$ | Tumor growth rate | $day^{-1}$ | 0.173 – 0.339 | [1,2] | 0 - 1 | 0.451 |
| $\boldsymbol{C_{max}}$ | Tumor carrying capacity | $mm^3$ | ~400 | [3] (Fig. 2A) | 100-500 | 375 |
| $\boldsymbol{k_{TC}}$ | T cell kill rate of tumor cells | $(day\ mm^3)^{-1}$ | 0.386 | [4], histology data | 0 – 1.5 | 0.509 |
| $\boldsymbol{k_{GC}}$ | Gem kill rate of tumor cells | $day^{-1}$ | -- | Flow cytometry data, [5] | 0 – 0.25 | 0.351 |
| $\boldsymbol{K_m}$ | Gem Michaelis constant | $\mu M$ | 160 | [6] | Fixed | 160 |
| **T CELLS ($T$)** | | | | | | |
| $\boldsymbol{n_{CT}}$ | Cancer-mediated T cell proliferation rate | $(day\ mm^3)^{-1}$ | <<2.079 | [7] | 0 – 0.5 | 0.229 |
| $\boldsymbol{s_{MT}}$ | MDSC suppression rate of T cells | $(day\ mm^3)^{-1}$ | 15.79 | [8], histology data | 0 - 20 | 5.4 |
| $\boldsymbol{d_T}$ | T cell natural death rate | $day^{-1}$ | 0.01 | [9] | Fixed* | 0.01 |
| $\boldsymbol{T_0}$ | Homeostatic T cell population | $mm^3$ | 0.75 | Histology data | 0 – 1 | 0.227 |
| $\boldsymbol{k_{GT}}$ | Gem kill rate of T cells | $day^{-1}$ | 0.66 | Flow cytometry data, [5] | 0 – 1 | 1.27 |
| **MYELOID-DERIVED SUPPRESSOR CELLS, MDSCS ($M$)** | | | | | | |
| $\boldsymbol{r_{CM}}$ | Cancer recruitment rate of MDSCs | $day^{-1}$ | 0.0880 – 0.176 | [10] | Fixed* | 0.132 |
| $\boldsymbol{s_{TM}}$ | T cell suppression of MDSCs | $(day\ mm^3)^{-1}$ | -- | Estimated | 0 - 5 | 0.0249 |
| $\boldsymbol{d_M}$ | MDSC natural death rate | $day^{-1}$ | 3.07 | [11] | Fixed* | 3.07 |
| $\boldsymbol{M_0}$ | Homeostatic MDSC population | $mm^3$ | 0.825 | Histology data | 0 - 1 | 0.0485 |
| $\boldsymbol{k_{GM}}$ | Gem kill rate of MDSCs | $day^{-1}$ | 0.46 | Flow cytometry data, [5] | 0 – 0.5 | 0.398 |
| **GEMCITABINE, GEM ($G$)** | | | | | | |
| $\boldsymbol{d_G}$ | Gem decay rate | $\mu M^{-1}$ | 1.54 | Data - private communication | Fixed | 1.54 |

*$r_{CM} \in [0,0.5]$ during virtual cohort generation, although all accepted virtual mice had $r_{CM}$ values between 0 and 0.3 $day^{-1}$. For sensitivity analysis, $r_{CM} \in [0,0.5]$, $\boldsymbol{d_T} \in [0,0.5]$, and $\boldsymbol{d_M} \in [0.25,5]$.

### S1.1 Treatment injections

#### Injection of Gem ($\boldsymbol{u_G}$)

Mice were treated with $500\ \mu g$ of Gem in $50\ \mu L$ PBS (phosphate buffer saline) [5]. Since we modelled the effect of Gem using Michaelis-Menten dynamics, we converted this to be in terms of concentration, $\mu M$. Gem has a molecular weight of $263.20\ g/mol$ [12]. Therefore, we calculated the concentration of the Gem injection to be:

$$u_G = \frac{500\ \mu g}{50\ \mu L} = \frac{10g}{1\ L} = \left(\frac{10g}{1\ L}\right)\left(\frac{1\ mol}{263.20\ g}\right)\left(\frac{1\ \mu\, mol}{10^{-6}\ mol}\right) = 3.80 \times 10^4\ \mu M.$$

#### Injection of OT-1 ($\boldsymbol{u_T}$)

From our histology image at day 17, the average area of a CD8$^+$ T cell is $5.2383 \times 10^{-5}\ mm^2$. Translating this area to a sphere, the average volume is $2.852 \times 10^{-7}\ mm^3$. Our injected OT-1 dose at day 14 was $5 \times 10^6$ cells [5], which converts to a volume of $1.426\ mm^3$. According to human data (private communication), the percent of tumor-infiltrating lymphocytes remaining after injection is $75\%$. Therefore, we assumed that the injection of OT-1 cells, $u_T$, is $1.0695\ mm^3$.

### S1.2 Bladder cancer parameter values

#### Tumor growth rate ($\boldsymbol{p_C}$)

Our previous experiments with orthotopically implanted MB49-OVA tumors in a mouse model showed exponential growth of tumor cells [5]. The in vivo doubling time of MB49 tumors is about 4 days[1]. Thus, we used the following equation:

$$\frac{dC}{dt} = p_C C,$$

and obtained an estimate for the tumor growth rate, $p_C$, of $0.173\ day^{-1}$.

White-Gilbertson et al. [2] saw a higher MB49 proliferation rate of up to $0.339\ day^{-1}$. From their experiments, the wet weight of excised tumors is up to $230mg$ 14 days after implanting 200,000 MB49 bladder cancer cells. Epithelial tumors are about $10^8$ cancer cells per $1g$ wet weight [13]. $230mg$ of tumor corresponds to approximately $2.3 \times 10^7$ MB49 bladder cancer cells. Assuming the equation for exponential growth again, we got a tumor growth rate, $p_C$, of $0.339\ day^{-1}$. While both experiments were for subcutaneous murine models of MB49, the results give an idea of the in vivo proliferation rate of the MB49 bladder cancer cell line, which we allowed to be within a range of $[0,1]\ day^{-1}$.

##### T cell kill rate of tumor cells ($k_{TC}$)

Kuznetsov et al. [4] used a mathematical model along with in vivo data to predict that the T cell kill rate of tumor cells was $1.101 \times 10^{-7}\ day^{-1}\ cell^{-1}$. The average volume of a CD8+ T cell is $2.852 \times 10^{-7}\ mm^3$ from our histology data (data not shown). Using this volume to convert the T cell kill rate to be in terms of $(day\ mm^3)^{-1}$, $k_{TC} = 0.386\ \ day^{-1}\ mm^3$. The range in Table S1: $0\ -\ 1.5\ \ (day\ mm^3)^{-1}$ accounts for a range of T cell diameters ($5 - 10\ \mu m$ compared to our $8.2\ \mu m$) in other literature [14].

##### Gem Michaelis constant ($K_m$)

The Gem Michaelis constant, $K_m$, is defined as the Gem concentration at which the transporter protein is at half its maximum uptake rate. Mackey et al. [6] determined different Michaelis constants for Gem depending on three nucleoside transporter proteins: CNT1 ($K_m = 24\mu M$), ENT1 ($K_m = 160\mu M$), and ENT2 ($K_m = 740\mu M$). From this, depending on the transporter protein used for cellular uptake, the Michaelis constant for Gem can be different.

However, we assumed that $K_m$ is the same for each cell type in our model, i.e., the same transporter protein is used for bladder cancer cells, T cells, and MDSCs. We justified this as follows: Matsumura et al. [15] found that higher expression of ENT1 is associated with longer survival for bladder cancer patients treated with Gem, so we assumed that ENT1 plays a major role in Gem uptake for bladder cancer cells. Further, although no literature was found on the uptake of Gem by naïve T cells or OT-1 cells, Klysz et al. [16] found that ENT1 was upregulated in CAR-T cells, while ENT2 was downregulated. No literature was found regarding the transporter protein used by MDSCs for Gem uptake, so for model simplification, MDSCs were assumed to use ENT1. Thus, we used ENT1's Michaelis constant of $160\mu M$ for each cell type.

#### S1.3 T Cell parameter values

##### Cancer-mediated proliferation of T cells ($n_{CT}$)

The in vivo doubling time for CD8+ T cells is 2-8 hours depending on the context: disease (2 hours [17]), OT-1 cells (4.5 hours [18]), or in general (6-8 hours [7]). Assuming that T cells are modelled exponentially, this doubling time translates to a growth rate of $2.079 - 8.318\ day^{-1}$. However, we modelled T cell expansion as $n_{CT}CT$ where $n_{CT}$ is in terms of $day^{-1}(mm^3)^{-1}$, so we assumed that $n_{CT}$ should be within a smaller range given the volume of the tumor. Dividing $8.318\ day^{-1}$ by the initial mean tumor volume at day 6 ($27.7\ mm^3$), the updated upper bound is $0.3\ day^{-1}(mm^3)^{-1}$. So, parameter estimation used the range of $0\ -\ 0.5\ day^{-1}(mm^3)^{-1}$.

#### MDSC suppression rate of T cells $(s_{MT})$

Anderson et al. [8] fitted a mathematical model to glioblastoma data and determined that the probability distribution mode for the MDSC suppression rate of T cells was $3.60 \times 10^{-6}\ day^{-1}\ cell^{-1}$. From our Ly6G histology data, the average area of a Ly6G-stained cell is $45.07\ \mu m^2$, which equates to a volume of $2.28 \times 10^{-7}\ mm^3$. Thus, the $s_{MT}$ estimate from [8] converts to $15.79\ day^{-1}(mm^3)^{-1}$.

#### Homeostatic T cell population $(T_0)$

We had two histology data points for a mouse bladder without an implanted tumor. In $1\ mm^3$ of tissue, mouse 1 had an average of $0.00519\ mm^3$ CD8⁺ T cells, while mouse 2 had an average of $7.72 \times 10^{-4}\ mm^3$ CD8⁺ T cells. So, we assumed that the homeostatic T cell population is at a density of $0.003\ mm^3$ of CD8⁺ T cells in $1\ mm^3$ of tissue. Since the mouse must be humanely euthanized once the tumor reaches $250\ mm^3$ in size, we took the homeostatic T cell population to be the volume of T cells within that volume in a non-cancerous mouse, thus $T_0 = \left(\frac{0.003\ mm^3\ of\ CD8^+\ T\ cells}{1\ mm^3\ of\ tissue}\right) 250\ mm^3 = 0.75\ mm^3$ of CD8⁺ T cells.

Since it is unlikely that all naïve T cells within the $250\ mm^3$ volume interact with a newly implanted tumor, this calculation was more for the purpose of providing a rough upper bound for the homeostatic T cell population. As our $T_0$ estimate showed in the S1 Table, the fitted value was lower than this calculation, which we expected.

#### Gem kill rate of T cells $(k_{GT})$

Using flow cytometry data at day 14 ([5], Fig 4), we roughly estimated of the Gem kill rate of T cells. We simplified the T cell equation and assumed that T cells are constant except under treatment with Gem:

$$\frac{dT}{dt} = -k_{GT} T \frac{G}{K_m + G}.$$

Since Gem decays exponentially, $G(t) = G(0)e^{-d_G t}$, so

$$T(t) = T(0)\ e^{-k_{GT}\left(\frac{G(t)}{K_m + G(t)}\right) t} = T(0)\ e^{-k_{GT}\left(\frac{G(0)e^{-d_G t}}{K_m + G(0)e^{-d_G t}}\right) t}.$$

Gem was treated at a concentration of $3.80 \times 10^4\ \mu M$ and the Gem decay rate, $d_G$, is calculated to be $1.54\ day^{-1}$ (see S1.1.1 and S1.5.1). The Michaelis constant, $K_m$, is estimated in [6] to be $160\ \mu M$. So, the population of T cells 4 days after Gem treatment is:

$$T_{gem}(4) = T_{gem}(0)\ e^{-k_{GT}\left(\frac{80.27}{160+80.27}\right) 4} = T_{gem}(0)\ e^{-k_{GT}(1.3364)}.$$

From flow cytometry data, the following ratio was taken 4 days after treatment:

$$\frac{average\ \#CD8\ /\ mg\ tumor\ in\ gem\ cohort}{average\ \#CD8\ /\ mg\ tumor\ in\ untx\ cohort} = 0.413.$$

Therefore, $T_{gem}(4) = T_{untx}(4) * 0.413$. Assuming the T cell population is constant without Gem treatment,

$$T_{gem}(4) = T_{untx}(4) * 0.413 = T_{gem}(0) * 0.413.$$

Altogether, this rearranged becomes

$$0.413 = \frac{T_{gem}(4)}{T_{gem}(0)} = e^{-k_{GT}(1.3364)},$$

implying that $k_{GT} = \frac{ln(0.413)}{-1.3364} = 0.66$.

#### S1.4 MDSC parameter values

##### Cancer recruitment of MDSCs ($r_{CM}$)

Zhang et al. [10] performed in vitro migration assays evaluating the migration of MDSCs due to the presence of J82 bladder cancer cells. $1 \times 10^6$ MDSCs were put in the upper migration chamber, while $5 \times 10^6$ J82 cells were placed in the lower chamber. Then, after waiting 12-24 hours for incubation, the number of migrated MDSCs were counted. For the control without the cancer cells, approximately 17% of MDSCs migrated to the lower chamber due to random migration. However, when J82 cells were present, about 61% of MDSCs migrated. This results in 44% of MDSCs migrating due to the presence of bladder cancer cells.

Using a simplified MDSC equation for recruitment,

$$\frac{dM}{dt} = r_{CM}C = r_{CM}\,(5 \times 10^6),$$

implying that the number of migrated MDSCs, $M$, to the lower chamber is

$$M = r_{CM}\,(5 \times 10^6)\,t\ .$$

As expected, at $t = 0$ days, there are $0$ migrated MDSCs. By $t = 0.5 - 1\ day$, $(1 \times 10^6)(44\%) = 4.4 \times 10^5$ MDSCs have migrated. Calculating for $r_{CM}$, we got

$$r_{CM} = \frac{4.4 \times 10^5}{(5 \times 10^6)t} \text{ for } t = 0.5 - 1\ \ days$$

which produced the range of $r_{CM} \in [0.0880{,}0.1760]\ day^{-1}$.

#### T cell suppression rate of MDSCs $(s_{TM})$

From day 17 histology data for the OT-1 treated versus non-OT-1 treated cohorts (Fig 3), we assumed that T cells had a role in decreasing the population of MDSCs. This assumption was confirmed through conversations with our experimentalists. Although no parameter value could be found from literature, it was reasonable to assume that this suppression is much less than the MDSC suppression of T cells $(s_{MT})$. Therefore, $s_{TM}$ was assumed to have a range of $0 - 5\ day^{-1}(mm^3)^{-1}$.

#### Homeostatic MDSC population $(M_0)$

Like the calculation for the homeostatic T cell population $(T_0)$, we used histology data from two mice not implanted with a tumor to estimate $M_0$. In the bladder, the density of MDSCs for mouse 1 was $0.00451\ mm^3$ MDSCs per $1\ mm^3$ of tissue, while mouse 2 exhibited $0.00214\ mm^3$ MDSCs per $1\ mm^3$ of tissue. Using the maximum allowable volume for the tumor size before euthanasia (i.e., $250\ mm^3$) to give an upperbound on the spatial spread of homeostatic MDSCs, we calculated that the homeostatic MDSC population to be $M_0 = \left(\frac{0.0033\ mm^3\ of\ MDSCs}{1\ mm^3\ of\ tissue}\right) 250\ mm^3 = 0.825\ mm^3$ MDSCs.

Since it is unlikely that all MDSCs within the $250\ mm^3$ volume interact with a newly implanted tumor, this calculation was more for the purpose of providing a rough upperbound estimate of the homeostatic MDSC population. As our $M_0$ estimate showed in Table S1, the fitted value was a lot lower than this calculation, which we expected.

#### MDSC natural death rate $(d_M)$

In vivo MDSCs have a lifespan of 1-2 days [11]. We assumed that, on average, the MDSC lifespan is 1.5 days. Using the following equation,

$$\frac{dM}{dt} = -d_M M,$$

and the assumption that $M(0) = 100\%$, while $M(1.5) = 1\%$, we obtain

$$d_M = \frac{ln(0.01)}{-1.5} = 3.07\ day^{-1}.$$

#### Gem kill rate of MDSCs $(k_{GM})$

Following the same assumption of our calculation for the Gem kill rate of T cells, $k_{GT}$, in S1.3.4, we used the ratio from flow cytometry 4 days after Gem treatment:

$$\frac{average\ \#MDSCs\ /\ mg\ tumor\ in\ gem\ cohort}{average\ \#MDSCs\ /\ mg\ tumor\ in\ untx\ cohort} = 0.543,$$

and calculated a rough estimate of the Gem kill rate of MDSCs:

$$k_{GM} = \frac{ln(0.543)}{-1.3364} = 0.46.$$

Since the Gem dose in experiments was chosen to not be lethal for cancer cells but to result in lymphodepletion of immunosuppressive cells, we assumed that $k_{GC} \leq \min(k_{GT}, k_{GM})$, so we rejected parameter sets that did not fulfill this condition during parameter fitting.

#### S1.5 Gem parameter values

##### Gem decay rate $(d_G)$

Since Gem is assumed to be cleared from the tumor tissue within 3 days after injection (private communication), we calculated the Gem decay rate as follows:

$$\frac{dG}{dt} = -d_G G,$$

and $G(0) = 100\%$, while $G(3) = 1\%$, we obtained

$$d_G = \frac{ln(0.01)}{-3} = 1.54\ day^{-1}.$$

### S2 Equilibrium and stability analysis

In this supplementary section, we determined the tumor-free equilibrium and its stability to ensure that the immune landscape of our model's tumor-free equilibrium is consistent with histology data from a normal tumor-free mouse bladder, which shows presence of $CD8^+$ T cells and $Ly6G^+$ MDSCs. Further, by considering the stability of the tumor-free equilibrium, we ascertained if our model implicitly assumes that recurrence is inevitable.

We found a unique biological tumor-free equilibrium of our model in Theorem A, and then in Theorem B, we determined its local asymptotic stability. For both theorems, we used the simpler version of the model without treatment:

$\boldsymbol{Cancer\ cells\ (C)}$:

$$\frac{dC}{dt} = \underbrace{p_C C\left(1 - \frac{C}{C_{max}}\right)}_{logistic\ growth} - \underbrace{k_{TC} TC}_{\substack{T\ cell \\ killing}}$$

$\boldsymbol{T\ cells\ (T)}$:

$$\frac{dT}{dt} = \underbrace{n_{CT} CT}_{\substack{cancer-mediated \\ proliferation}} - \underbrace{s_{MT} MT}_{\substack{MDSC \\ suppression}} - \underbrace{d_T T}_{\substack{natural \\ death}} + \underbrace{h_T}_{\substack{homeostatic \\ influx}}$$

$\boldsymbol{Myeloid - derived\ suppressor\ cells\ (MDSCs)\ (M)}$:

$$\frac{dM}{dt} = \underbrace{r_{CM} C}_{\substack{MDSC \\ recruitment}} - \underbrace{s_{TM} TM}_{\substack{T\ cell \\ suppression}} - \underbrace{d_M M}_{\substack{natural \\ death}} + \underbrace{h_M}_{\substack{homeostatic \\ influx}}$$

**Theorem A. Tumor-free equilibria.** The system has a unique biological tumor-free equilibrium, $(C, T, M) = (0, T_+^*, M_+^*)$, in quadrant 1, and a unique non-biological tumor-free equilibrium, $(C, T, M) = (0, T_-^*, M_-^*)$, in quadrant 3, where $T_\pm^* = \frac{h_T}{s_{MT} M_\pm^* + d_T}$ , $M_\pm^* = \frac{-\beta \pm \sqrt[2]{\beta^2 + 4 s_{MT} d_M d_T h_M}}{-2 s_{MT} d_M}$ , and $\beta = h_M s_{MT} - s_{TM} h_T - d_T d_M$.

**Proof.** Assume $C = 0$. Setting $\frac{dT}{dt} = 0$, we get $0 = -s_{MT} MT - d_T T + h_T$. Solving for $T$,

$T = \frac{h_T}{s_{MT} M + d_T} > 0$ if $M \geq 0$.

Now, setting $\frac{dM}{dt} = 0$, we get

$$0 = -s_{TM}TM - d_M M + h_M.$$

Inserting $T$, we get

$$0 = -s_{TM}\left(\frac{h_T}{s_{MT}M + d_T}\right)M - d_M M + h_M.$$

Multiplying by $s_{MT}M + d_T$, we get

$$0 = -s_{TM}h_T M + (s_{MT}M + d_T)(-d_M M + h_M).$$

Rearranging

$$0 = -s_{MT}d_M M^2 + (h_M s_{MT} - s_{TM}h_T - d_T d_M)M + d_T h_M$$

Using the quadratic formula,

$$M = \frac{-\beta \pm \sqrt[2]{\beta^2 + 4s_{MT}d_M d_T h_M}}{-2s_{MT}d_M}$$

Where $\beta = h_M s_{MT} - s_{TM}h_T - d_T d_M$.

For $M_-$,

$$M_- = \frac{-\beta + \sqrt[2]{\beta^2 + 4s_{MT}d_M d_T h_M}}{-2s_{MT}d_M} = \frac{\beta - \sqrt[2]{\beta^2 + 4s_{MT}d_M d_T h_M}}{2s_{MT}d_M} < \frac{\beta - \sqrt[2]{\beta^2}}{2s_{MT}d_M} \leq 0.$$

For $M_+$,

$$M_+ = \frac{-\beta - \sqrt[2]{\beta^2 + 4s_{MT}d_M d_T h_M}}{-2s_{MT}d_M} = \frac{\beta + \sqrt[2]{\beta^2 + 4s_{MT}d_M d_T h_M}}{2s_{MT}d_M} > \frac{\beta + \sqrt[2]{\beta^2}}{2s_{MT}d_M} \geq 0.$$

So, $T_+ = \frac{h_T}{s_{MT}M_+ + d_T} > 0$. Thus, $(0, T_+^*, M_+^*)$ is an equilibrium point in the first quadrant where $T_+^*$ and $M_+^*$ are strictly positive.

Note that the equations $0 = -s_{MT}MT - d_T T + h_T$ and $0 = -s_{TM}TM - d_M M + h_M$ are symmetric, so if we solve for $T$ in the same way as for $M$, we get $T_-^* < 0$ and $T_+^* > 0$. Since $T_+^*$ corresponds to $M_+^*$, $T_-^*$ must correspond to $M_-^*$. So, there is a non-biological equilibrium point in the third quadrant, $(0, T_-^*, M_-^*)$, where $T_-^*$ and $M_-^*$ are strictly negative.

■

From the histology data, there were T cells and MDSCs when no tumor is present, so the biological tumor-free equilibrium aligns with the data. Since we did not have parameter values from literature on the homeostatic T cell, $h_T$, and MDSC, $h_M$, proliferation rate, we used the results from Theorem A to rewrite these parameters in terms of the tumor-free homeostatic T cell, $T_0$, and MDSC, $M_0$, population, for which we had histology data. From Theorem A's results, $h_T = (s_{MT}M_0 + d_T)T_0$. The more complicated equational form of $M_0$ makes it very difficult—if not impossible—to solve for $h_M$. However, using the simpler equational form found in the last paragraph of Theorem A, $h_M = (s_{TM}T_0 + d_M)M_0$.

In Theorem B, we determined conditions for the local asymptotic stability of the tumor-free equilibria from Theorem A. Local asymptotic stability of the biological tumor-free equilibrium means that if the tumor is sufficiently small, the system will continue to tend towards this equilibrium. Thus, if treatment diminishes the tumor size enough and if a mouse fulfills parametric conditions for local asymptotic stability, the mouse will continue to disease-free survival.

**Theorem B. Stability of the tumor-free equilibria.** The tumor-free equilibria, $(0, T_{\pm}{}^*, M_{\pm}{}^*)$, are locally asymptotically stable when $p_C - k_{TC}T_{\pm}{}^* < 0$, and saddle points when $p_C - k_{TC}T_{\pm}{}^* > 0$. This implies that the non-biological tumor-free equilibrium, $(0, T_-^*, M_-^*)$, is always a saddle point, but $(0, T_+^*, M_+^*)$ may be a saddle point or may be locally asymptotically stable.

**Proof.** The Jacobian at $C = 0$ is

$$\begin{pmatrix} p_C - k_{TC}T^* & 0 & 0 \\ n_{CT}T^* & -s_{MT}M^* - d_T & -s_{MT}T^* \\ r_{CM} & -s_{TM}M^* & -s_{TM}T^* - d_M \end{pmatrix},$$

Where the first eigenvalue is

$$\lambda_1 = p_C - k_{TC}T^*.$$

Solving for the other eigenvalues, we have

$$0 = (-s_{MT}M^* - d_T - \lambda)(-s_{TM}T^* - d_M - \lambda) - (s_{TM}M^*)(s_{MT}T^*)$$

Which simplifies to

$$0 = \lambda^2 + \big((s_{MT}M^* + d_T) + (s_{TM}T^* + d_M)\big)\lambda + (s_{MT}M^* + d_T)(s_{TM}T^* + d_M) - s_{TM}s_{MT}M^*T^*$$

Rewriting this, we have

$$0 \ = \ \lambda^2 + (a + b)\lambda + ab - c$$

Where

$$a = s_{MT}M^* + d_T$$

$$b = s_{TM}T^* + d_M$$

$$c = s_{TM}s_{MT}M^*T^*$$

By the quadratic formula,

$$\lambda = \frac{-(a+b) \pm \sqrt[2]{(a+b)^2 - 4(ab-c)}}{2}.$$

**Case 1.** Assume that $(a+b)^2 - 4(ab-c) \leq 0$.

Then, $Re(\lambda_2)$ and $Re(\lambda_3) < 0$, so the tumor-free equilibrium is locally asymptotically stable under the condition that $\lambda_1 = p_C - k_{TC}T^* < 0$.

**Case 2.** Assume $(a+b)^2 - 4(ab-c) > 0$.

For $\lambda_2$,

$$\lambda_2 = \frac{-(a+b) - \sqrt[2]{(a+b)^2 - 4(ab-c)}}{2} < 0.$$

For $\lambda_3$, note that, according to our case 2 assumption, $(a+b)^2 > 4(ab-c)$.

Note that

$$ab - c = (s_{MT}M^* + d_T)(s_{TM}T^* + d_M) - s_{TM}s_{MT}M^*T^* = d_Ts_{TM}T^* + d_Ms_{MT}M^* + d_Md_T > 0$$

So,

$$\lambda_3 = \frac{-(a+b) + \sqrt[2]{(a+b)^2 - 4(ab-c)}}{2} < \frac{-(a+b) + \sqrt[2]{(a+b)^2}}{2} = 0$$

So, $\lambda_2$ and $\lambda_3 < 0$. Thus, in both cases, the tumor-free equilibrium is locally asymptotically stable as long as $p_C - k_{TC}T^* < 0$.

■

Under this condition, the non-biological tumor-free equilibrium, $(0, T_-^*, M_-^*)$, is always a saddle point, but the biological tumor-free equilibrium, $(0, T_+^*, M_+^*)$, is locally stable if the immune response $(k_{TC}T^*)$ is greater than the tumor growth $(p_C)$. This implies that if a treatment regimen can get the tumor within a sufficient distance of the tumor-free equilibrium, a healthy individual would see permanent remission. According to our parameter estimates for an average mouse (Section 5), the local asymptotic stability condition was not fulfilled, so the model predicted that an average mice would experience recurrence after treatment, even if treatment can get the tumor quite small.

Different model formulations would assume that complete remission is essentially impossible by preventing the possibility of a locally asymptotically tumor-free equilibrium. Theorems C - D give an example of this situation by showing that if one removes the terms for homeostatic T cell $(h_T)$ and MDSC $(h_M)$ proliferation, the tumor-free equilibrium would be $(0,0,0)$ and that this equilibrium would be a saddle point. This equilibrium would firstly be inconsistent with our histology data from a normal tumor-free mouse bladder. Secondly, it would mean that unless the treatment regimen happened to get the tumor to the system's stable manifold, the model predicts that no mouse will progress to complete disease-free survival. While this may be the case, by providing the option for complete response, we allowed individual immune parameters to determine the chance of recurrence instead of creating a model that automatically assumes tumor recurrence.

For Theorems C - D, we used the following version of the treatment-free model, which excludes the homeostatic T cell $(h_T)$ and MDSC $(h_M)$ influx rate, to prove the importance of these terms for data fidelity and model assumptions.

$\boldsymbol{Cancer\ cells\ (C)}$:

$$\frac{dC}{dt} = \underbrace{p_C C\left(1 - \frac{C}{C_{max}}\right)}_{logistic\ growth} - \underbrace{k_{TC}TC}_{\substack{T\ cell\\ killing}}$$

$\boldsymbol{T\ cells\ (T)}$:

$$\frac{dT}{dt} = \underbrace{n_{CT}CT}_{\substack{cancer-mediated\\ proliferation}} - \underbrace{s_{MT}MT}_{\substack{MDSC\\ suppression}} - \underbrace{d_T T}_{\substack{natural\\ death}}$$

$\boldsymbol{Myeloid-derived\ suppressor\ cells\ (MDSCs)\ (M)}$:

$$\frac{dM}{dt} = \underbrace{r_{CM}C}_{\substack{MDSC\\ recruitment}} - \underbrace{s_{TM}TM}_{\substack{T\ cell\\ suppression}} - \underbrace{d_M M}_{\substack{natural\\ death}}$$

**Theorem C. Tumor-free equilibrium of alternative model.** The treatment-free model has a unique biological tumor-free equilibrium, $(0,0,0)$.

**Proof.** Assume $C = 0$. Automatically, $\frac{dC}{dt} = 0$.

Since the system is assumed to be in equilibrium, if $\frac{dT}{dt} = 0$, this implies

$$0 = -s_{MT}MT - d_T T.$$

Therefore, either $M = \frac{-d_T}{s_{MT}}$ or $T = 0$.

Setting $\frac{dM}{dt} = 0$, we get

$$0 = -s_{TM}TM - d_M M,$$

which corresponds to either $T = \frac{-d_M}{s_{TM}}$ or $M = 0$.

We conclude that there are two possible tumor-free equilibria, $(0,0,0)$ and $(0, \frac{-d_M}{s_{TM}}, \frac{-d_T}{s_{MT}})$, but only one of which is biologically reasonable. Thus, the treatment-free model has a unique biological tumor-free equilibrium, $(0,0,0)$.

■

**Theorem D. Stability of the tumor-free equilibrium for alternative model.** The unique tumor-free biological equilibrium, $(0,0,0)$, for the treatment-free model is a saddle point.

**Proof.** The Jacobian evaluated at the tumor-free equilibrium, $(0,0,0)$, is

$$\begin{pmatrix} p_C & 0 & 0 \\ 0 & -d_T & 0 \\ r_{CM} & 0 & -d_M \end{pmatrix},$$

which has eigenvalues $p_T > 0, -d_T < 0, -d_M < 0$, so the tumor-free equilibrium is a saddle point. This means that if we stop treatment before the tumor is exactly 0, tumor recurrence is inevitable.

■

### S3 Global sensitivity analysis figures at alternative times

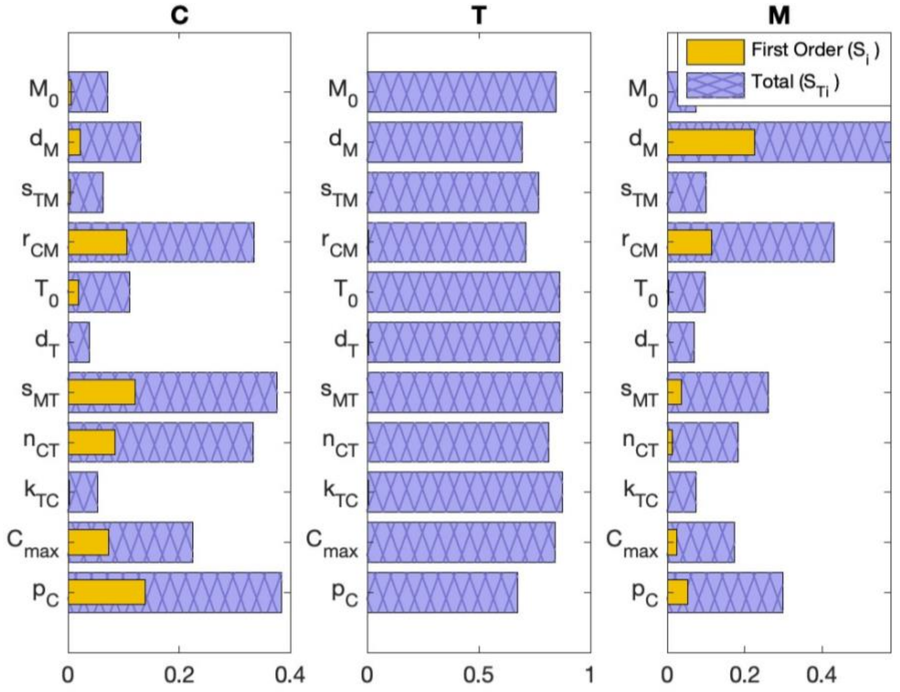


(a) day 10

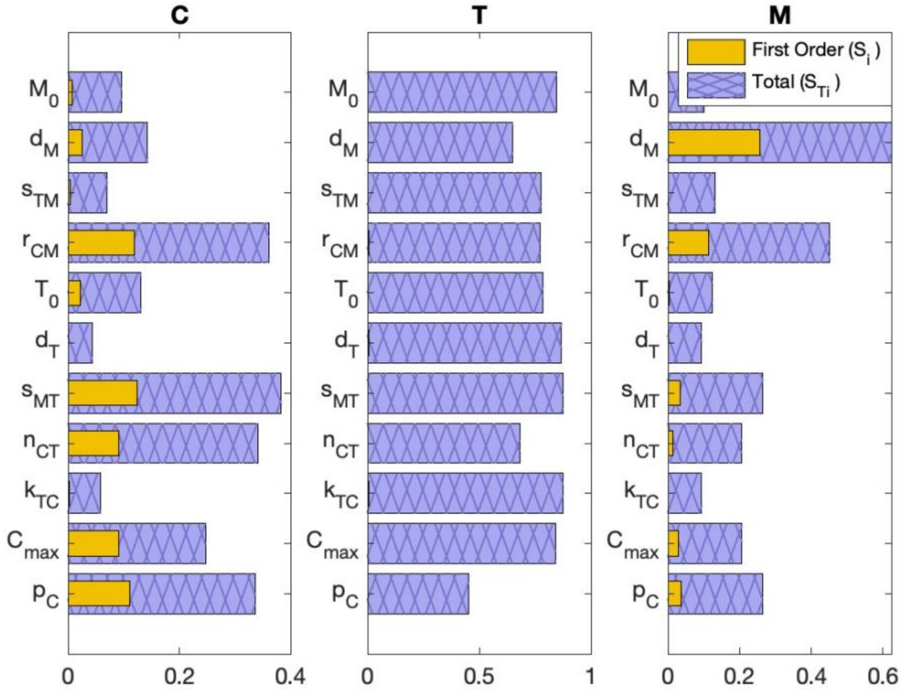


(b) day 15

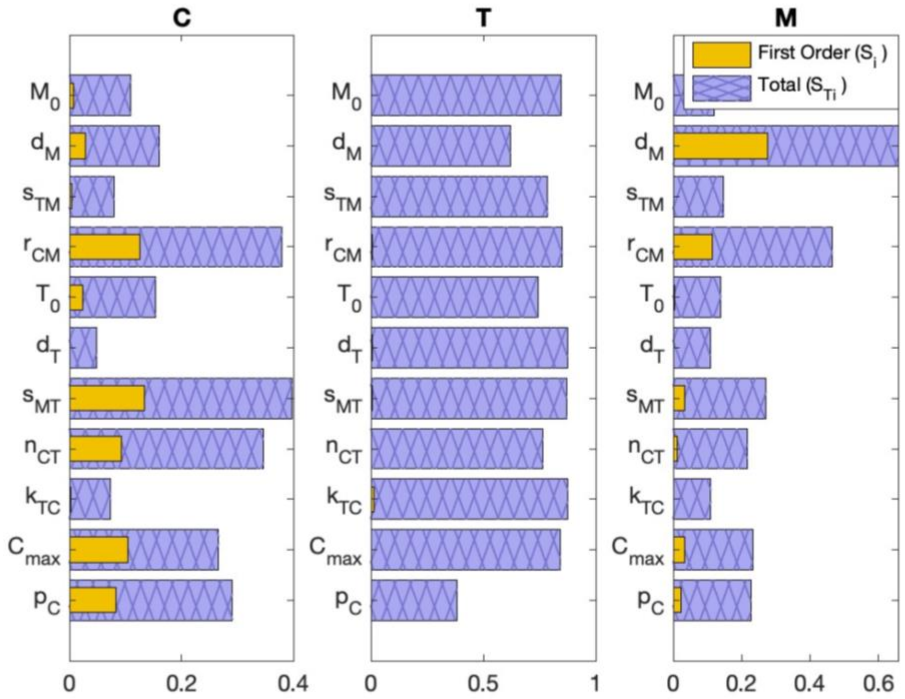


(c) day 25

**Fig S1. Global sensitivity analysis at alternative times.** eFAST was used to determine the treatment-free model's sensitivity to parameters after tumor-implantation on day 0. In Section 7 of the main text, we considered the treatment-free model's sensitivity to parameters at day 20 as this time point shows the most variability in data (Fig 3). Here, we evaluated additional time points during the treatment period (day (a) 10 and (b) 15) and slightly beyond ((c) day 25).

### References


1 Trachet, E. *MB49 - a bladder cancer murine tumor model*, <https://oncology.labcorp.com/mb49-bladder-cancer-murine-tumor-model> (2019).

2 White-Gilbertson, S., Davis, M., Voelkel-Johnson, C. & Kasman, L. M. Sex differences in the MB49 syngeneic, murine model of bladder cancer. *Bladder* **3**, e22 (2016).

3 Bunch, B. L. *et al.* Systemic and intravesical adoptive cell therapy of tumor-reactive T cells can decrease bladder tumor growth in vivo. *Journal for Immunotherapy of Cancer* **8**, e001673 (2020).

4 Kuznetsov, V. A., Makalkin, I. A., Taylor, M. A. & Perelson, A. S. Nonlinear dynamics of immunogenic tumors: parameter estimation and global bifurcation analysis. *Bulletin of mathematical biology* **56**, 295-321 (1994).

5 Bazargan, S. *et al.* Targeting myeloid-derived suppressor cells with gemcitabine to enhance efficacy of adoptive cell therapy in bladder cancer. *Frontiers in Immunology* **14**, 1275375 (2023).

6 Mackey, J. R. *et al.* Gemcitabine transport in xenopus oocytes expressing recombinant plasma membrane mammalian nucleoside transporters. *J Natl Cancer Inst* **91**, 1876-1881 (1999). https://doi.org:10.1093/jnci/91.21.1876

7 Lewis, D. A. & Ly, T. Cell cycle entry control in naïve and memory CD8+ T cells. *Frontiers in Cell and Developmental Biology* **9**, 727441 (2021).

8 Anderson, H. G. *et al.* Global stability and parameter analysis reinforce therapeutic targets of PD-L1-PD-1 and MDSCs for glioblastoma. *J Math Biol* **88**, 10 (2023). https://doi.org:10.1007/s00285-023-02027-y

9 Ribeiro, R. M., Mohri, H., Ho, D. D. & Perelson, A. S. In vivo dynamics of T cell activation, proliferation, and death in HIV-1 infection: why are CD4+ but not CD8+ T cells depleted? *Proceedings of the National Academy of Sciences* **99**, 15572-15577 (2002).

10 Zhang, H. *et al.* CXCL2/MIF-CXCR2 signaling promotes the recruitment of myeloid-derived suppressor cells and is correlated with prognosis in bladder cancer. *Oncogene* **36**, 2095-2104 (2017).

11 Ostrand-Rosenberg, S. & Fenselau, C. Myeloid-derived suppressor cells: immune-suppressive cells that impair antitumor immunity and are sculpted by their environment. *The Journal of Immunology* **200**, 422-431 (2018).

12 Information, N. C. f. B. PubChem Compound Summary for CID 60750, Gemcitabine. (2025).

13 Del Monte, U. Does the cell number 109 still really fit one gram of tumor tissue? *Cell cycle* **8**, 505-506 (2009).

14 Jiang, X. *et al.* MRI of tumor T cell infiltration in response to checkpoint inhibitor therapy. *Journal for immunotherapy of cancer* **8**, e000328 (2020).

15 Matsumura, N. *et al.* The prognostic significance of human equilibrative nucleoside transporter 1 expression in patients with metastatic bladder cancer treated with gemcitabine-cisplatin-based combination chemotherapy. *BJU Int* **108**, E110-116 (2011). https://doi.org:10.1111/j.1464-410X.2010.09932.x

16 Klysz, D. D. *et al.* Inosine induces stemness features in CAR-T cells and enhances potency. *Cancer Cell* **42**, 266-282 e268 (2024). https://doi.org:10.1016/j.ccell.2024.01.002
17 Yoon, H., Kim, T. S. & Braciale, T. J. The cell cycle time of CD8+ T cells responding in vivo is controlled by the type of antigenic stimulus. *PloS one* **5**, e15423 (2010).
18 Kurts, C., Kosaka, H., Carbone, F. R., Miller, J. F. & Heath, W. R. Class I–restricted cross-presentation of exogenous self-antigens leads to deletion of autoreactive CD8+ T cells. *The Journal of experimental medicine* **186**, 239-245 (1997).
